# Novel target identification using single cell sequencing and electrophysiology of cardiac sympathetic neurons in disease

**DOI:** 10.1101/2019.12.11.872580

**Authors:** Harvey Davis, Neil Herring, David J Paterson

**Affiliations:** Burdon Sanderson Cardiac Science Centre, Department of Physiology, Anatomy and Genetics, Sherrington Building, University of Oxford, Oxford, OX1 3PT, UK; Wellcome Trust OXION Initiative in Ion Channels and Disease, Department of Physiology, Anatomy and Genetics, Sherrington Building, University of Oxford, Oxford, OX1 3PT, UK; Oxford Heart Centre, Oxford University Hospitals NHS Foundation Trust, John Radcliffe Hospital, Oxford, OX3 9DU, UK

**Keywords:** Stellate Ganglia, Sympathetic, Dysautonomia, Hypertension, M-current

## Abstract

The activity of cardiac sympathetic nerves from the stellate ganglia is increased in many cardiovascular diseases contributing to the pathophysiology, however the mechanisms underlying this are unknown. Moreover, clinical studies show their surgical removal is an effective treatment, despite the biophysical properties of these neurons being largely unstudied. Here we demonstrate that stellate ganglia neurons from prehypertensive spontaneously hypertensive rats are hyperactive and describe in detail their electrophysiological phenotype guided by single cell RNA-sequencing, molecular biology and perforated patch-clamp to uncover the underlying mechanism. The expression of key transcripts was confirmed in human stellate ganglia. We further demonstrate the contribution of a plethora of ion channels to stellate ganglia neuronal firing, and show that hyperexcitability was curbed by M-current activators, non-selective sodium current blockers or inhibition of Na_v_1.1-1.3, Na_v_1.6 or I_NaP_. These findings have implications for target discovery to reduce cardiac sympathetic activity without resorting to surgery.

## Introduction

The stellate ganglia, which contain the cell bodies of sympathetic neurons that predominantly innervate the heart (Rajendran *et al*., 2019), are of considerable interest to cardiovascular research (Herring, Kalla and Paterson, 2019). Surgical removal of these ganglia is being used with great success clinically to treat life threatening ventricular arrhythmias associated with structural heart disease and some inherited arrhythmia syndromes (Schwartz, 2014; Herring, Kalla and Paterson, 2019). Despite this, the electrophysiology of these neurons remains largely unstudied.

Sympathetic hyperactivity is known to contribute to the pathophysiology of a variety of cardiovascular diseases, including hypertension, myocardial infarction, and heart failure, although the cellular mechanisms responsible for this have not been elucidated (Herring, Kalla and Paterson, 2019). In hypertension, increased cardiac sympathetic drive is linked to the development of left ventricular hypertrophy (Schlaich Markus, Kaye David, Lambert Elisabeth, Sommerville Marcus, Socratous Flora, 2003) and subsequent heart failure, which are independent predictors of mortality (Levy *et al*., 1990). A significant component of this sympathetic hyperactivity resides at the level of the post-ganglionic sympathetic neuron. For example, neurons from the stellate ganglia, have enhanced Ca^2+^ driven (Larsen *et al*., 2016) norepinephrine release (Shanks, Manou-Stathopoulou, *et al*., 2013) and impaired re-uptake via the NET transporter (Shanks, Mane, *et al*., 2013). However, both animal models and patients with hypertension also have increased sympathetic nerve firing rate as measured by muscle and renal sympathetic nerve activity (Grassi, 2009). Whether this is centrally driven or results from changes in the excitability of post-ganglionic neurons before the onset of hypertension is unknown (Sverrisdóttir *et al*., 2014), as are the ion channels involved in the sympathetic phenotype.

In this study we provide single cell RNA sequencing (scRNAseq) analysis of stellate ganglia neurons and use this to guide a detailed biophysical characterisation in an animal model of prehypertension (Minami *et al*., 1989), with validation in human stellate ganglia tissue. We highlight several important ion channels as putative targets for pharmacological intervention to reduce cardiac sympathetic neuron firing rate, rather than resorting to irreversible surgery with its associated complications (such as pneumothorax and Horner’s syndrome (Vaseghi *et al*., 2017)).

## Results

### Stellate ganglia neurons are hyperexcitable in the prehypertensive SHR

Perforated patch clamp measurements demonstrate that stellate ganglia neurons from prehypertensive spontaneously hypertensive rats (SHRs) have a significantly higher firing rate than neurons from normotensive age matched Wistars as shown in Figures 1A and 1B. This firing rate difference appears to be time resolved, with the majority of Wistar neurons firing action potentials within only the first 300 ms of stimulation. This phenotype is represented in Figure 1C via a raster plot of 30 Wistar and SHR neurons during a 1000 ms of 150 pA current injection. We also observed other indicators of cellular hyperexcitability. The change in firing rate was accompanied by a 3.1 ± 1.1 mV decrease in resting membrane potential between Wistar and SHR neurons (Figure 1D). The rheobase (minimum current injection of duration >300 ms required to reach the action potential threshold) was decreased in SHR neurons as observed via a series of 10 pA current steps of duration 1000 ms in the range 0-200 pA in amplitude (Figure 1E).

**Figure 1.**
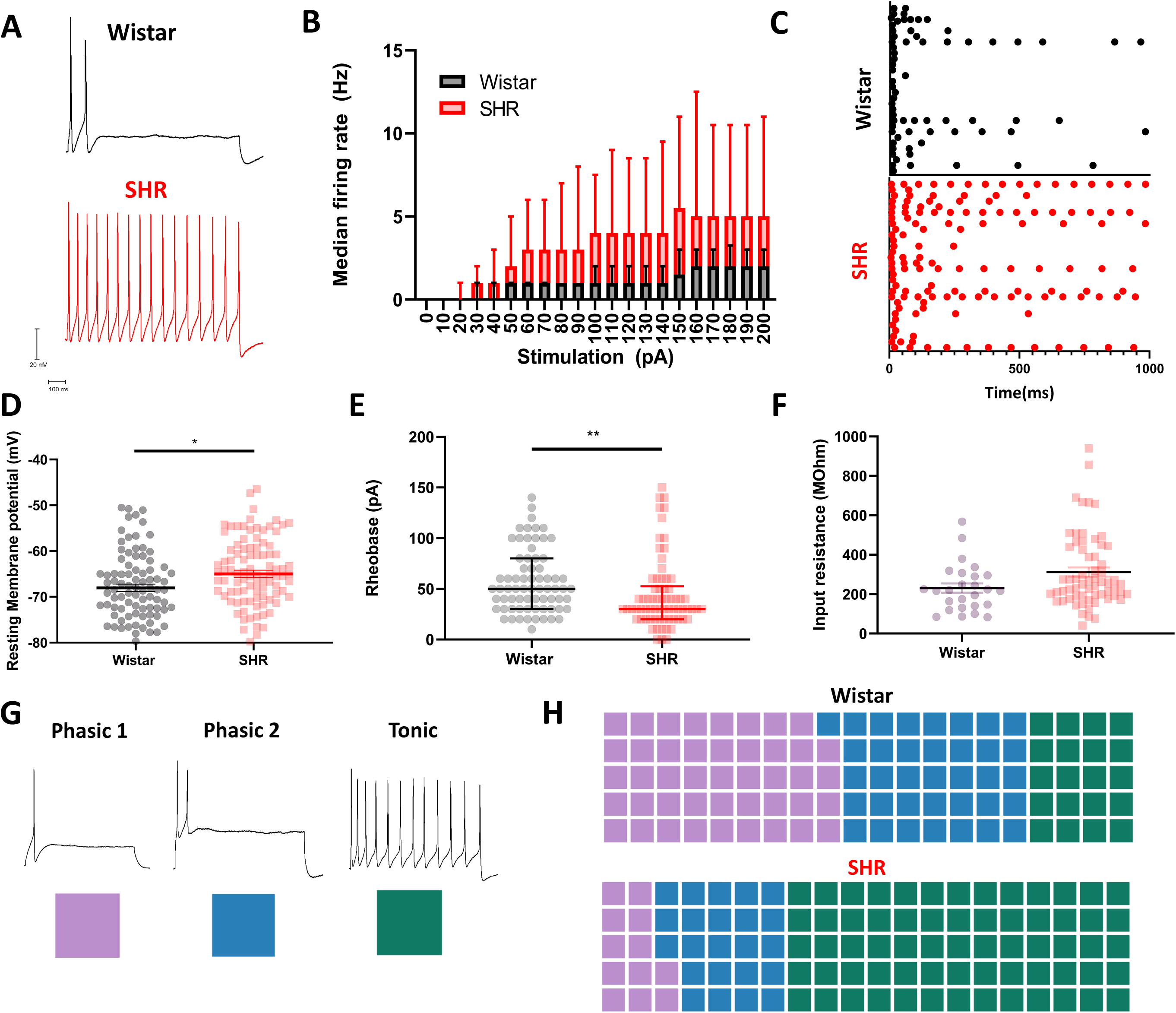
Stellate ganglia neurons of the spontaneously hypertensive rat have a hyperactive phenotype with an altered electrophysiological profile as observed by perforated patch clamp recordings. (A) Example traces showing the response of sympathetic neurons from control Wistar and pre-hypertensive SHR rats to 150 pA of current injection for a duration of 1000 ms. (B) Median response of Wistar (Black) and SHR (Red) neurons to a range of current injections (Wistar, n = 66; SHR, n = 69) (Mixed-effects model, p < 0.0001). (C) Time course of induced firing following a 1000 ms 150 pA current injection in 30 example Wistar and SHR neurons. (D) A significant positive shift in the resting membrane potential was observed in SHR neurons (Mean ± SEM) (Wistar, −68.04 ± 0.81 mV, n = 89; SHR, −64.97 ± 0.78 mV, n = 97) (Unpaired t-test, p = 0.0072). (E) A significant decrease in rheobase was observed in SHR neurons (Median ± IQR) (Wistar, 50 pA, n = 70; SHR, 30 pA, n = 70) (Mann-Whitney test, p = 0.0013). (F) A non-significant trend to increased input resistance was observed between Wistar and SHR neurons (Median ± IQR) (Wistar, 218.2 MΩ, n = 26; SHR, 251.8 MΩ, n = 63) (Mann Whitney test, p = 0.066). (G) Examples of sympathetic neuron firing rates are identified by their classical nomenclature. Here the colour scheme used to indicate these subtypes throughout the article are indicated by a colored square. (H) A trend towards tonic firing neurons was observed in the SHR, shown as percentage of cells conforming to each subtype (Wistar, n = 66; Phasic 1 n = 29, Phasic 2 n = 24, Tonic n = 13; SHR, n = 73; Phasic 1 n = 9, Phasic 2 n = 17, Tonic n = 47) (χ^2^ = 12.37, df =2, p-value = 0.0021).

### The SHR has a higher percentage of tonic firing neurons

Neuronal firing behaviour has been previously characterised into 3 subtypes (Cassell, Clark and McLachlan, 1986). Phasic 1 neurons, represent neurons that fire only one action potential for a 1000 ms current injection in the range of 0-200 pA. Phasic 2 neurons fire 2-5 action potentials, all within the first 500 ms of a 1000 ms stimulation pulse for all recordings within the stimulation range 0-200 pA. Tonic neurons fire either greater than 6 action potentials or continue to fire after 500 ms of stimulation of a 1000 ms stimulation pulse within the current injection range 0-200 pA. In Figure 1H and subsequent figures these subtypes have been assigned colour codes to allow for a visual representation of the firing rate and time course of firing. When the percentage of neurons conforming to these subtypes is compared between Wistar and SHR neurons, Wistar neurons were found to be predominantly phasic 1 and phasic 2, whereas in the SHR, neurons were predominantly found to be of the tonic subtype. These data are supported by similar observations in whole cell patch-clamp recordings (Supplementary Figure 1).

### A change in ion channel subunit expression was observed in SHR neurons

In order to investigate the underlying ion channel differences responsible for the difference in firing rate between SHR and Wistar stellate neurons, we undertook a scRNAseq analysis of the ganglia. ScRNAseq revealed a heterogenous population of cell clusters in the both Wistar and SHR dissociated stellate ganglia (Figure 2A), which mapped to known cell type markers (Figure 2B; Supplementary Figures 4 and 5) including two distinct population of cells which are specific for the full range of known sympathetic markers (Figure 2C; Supplementary Figure 6). The main difference between these two sympathetic neuronal populations is that one also expressed the co-transmitter neuropeptide-Y. Differential expression analysis between the Wistar and SHR sympathetic neuron populations identified in Figure 2C highlight a significant decrease in 5 ion channel subunit encoding genes that may contribute to firing rate of stellate ganglia sympathetic neurons (Table 1) (Full list in supplementary 2D).

**Table 1.**
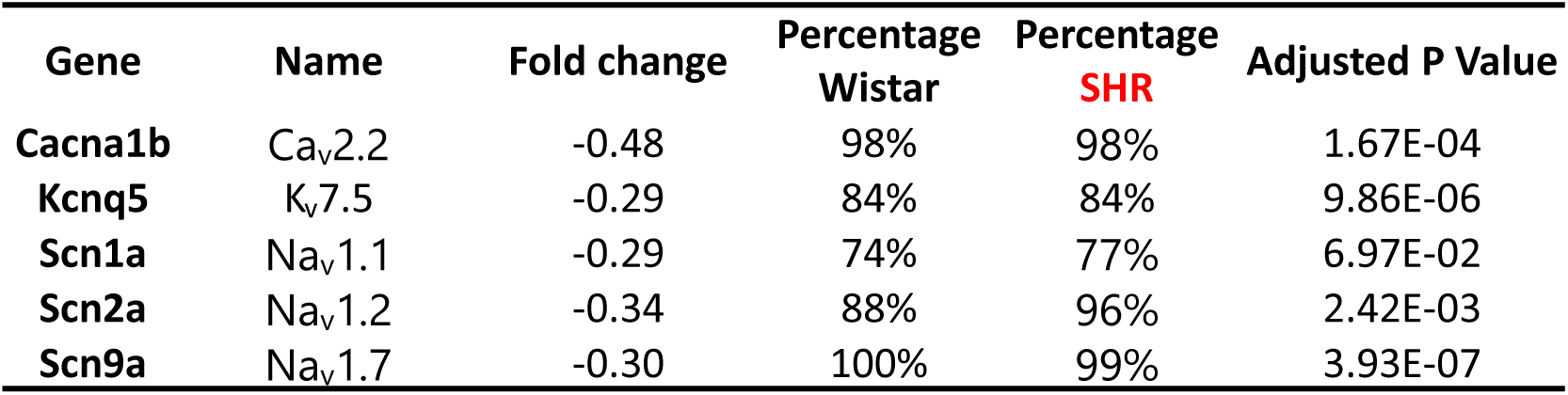
Differential expression analysis of key ion channels in Wistar vs SHR neurons reveals a number of ion channel subunits are downregulated at transcriptional level in the SHR. Differential expression analysis was performed using MAST in the Seurat package.

**Figure 2.**
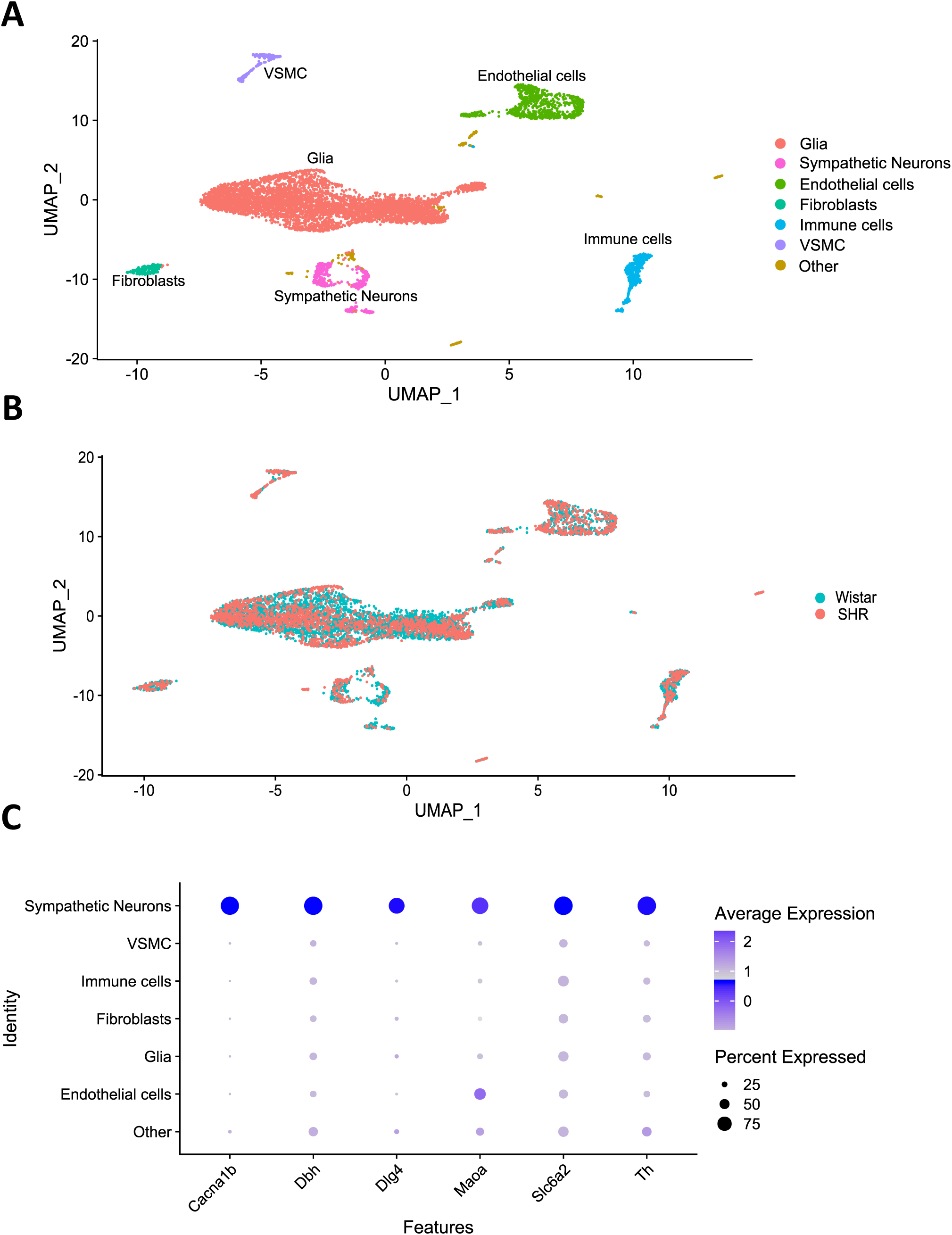
Single cell RNA-sequencing on Wistar and SHR populations reveals altered channel subunit expression in SHR stellate ganglia neurons. (A) Single cell RNA-sequencing reveals multiple clusters of cell transcriptomes. (B) These cell clusters align to multiple cell types, as defined using the best-known markers for these cell types. (C) Markers used for sympathetic neurons are shown to be primarily expressed in sympathetic neurons (Cacna1b = Ca_v_2.2; Dbh = Dopamine β hydroxylase; Dlg4 = PSD-95; Maoa = Monoamine oxidase A; Slc6a2 = NET; TH = Tyrosine hydroxylase).

### M-current is functionally reduced in SHR stellate ganglia neurons and present in human stellate ganglia

Was a decrease in KCNQ5 subunit expression the most likely explanation for the difference in phenotype? When assessed by RT-qPCR, gene expression of M-current encoding KCNQ2, KCNQ3 and KCNQ5 subunits is decreased in total RNA extracted from whole SHR ganglia (Figure 3A). We also confirmed M-current subunit expression in samples of total RNA taken from human stellate ganglia (Figure 3B). Using immunohistochemistry, M-current encoding subunits KCNQ2, KCNQ3 and KCNQ5, were also shown to be expressed at a protein level in tyrosine hydroxylase (TH) positive cells (Supplementary Figure 7G), a classic marker for sympathetic neurons.

**Figure 3.**
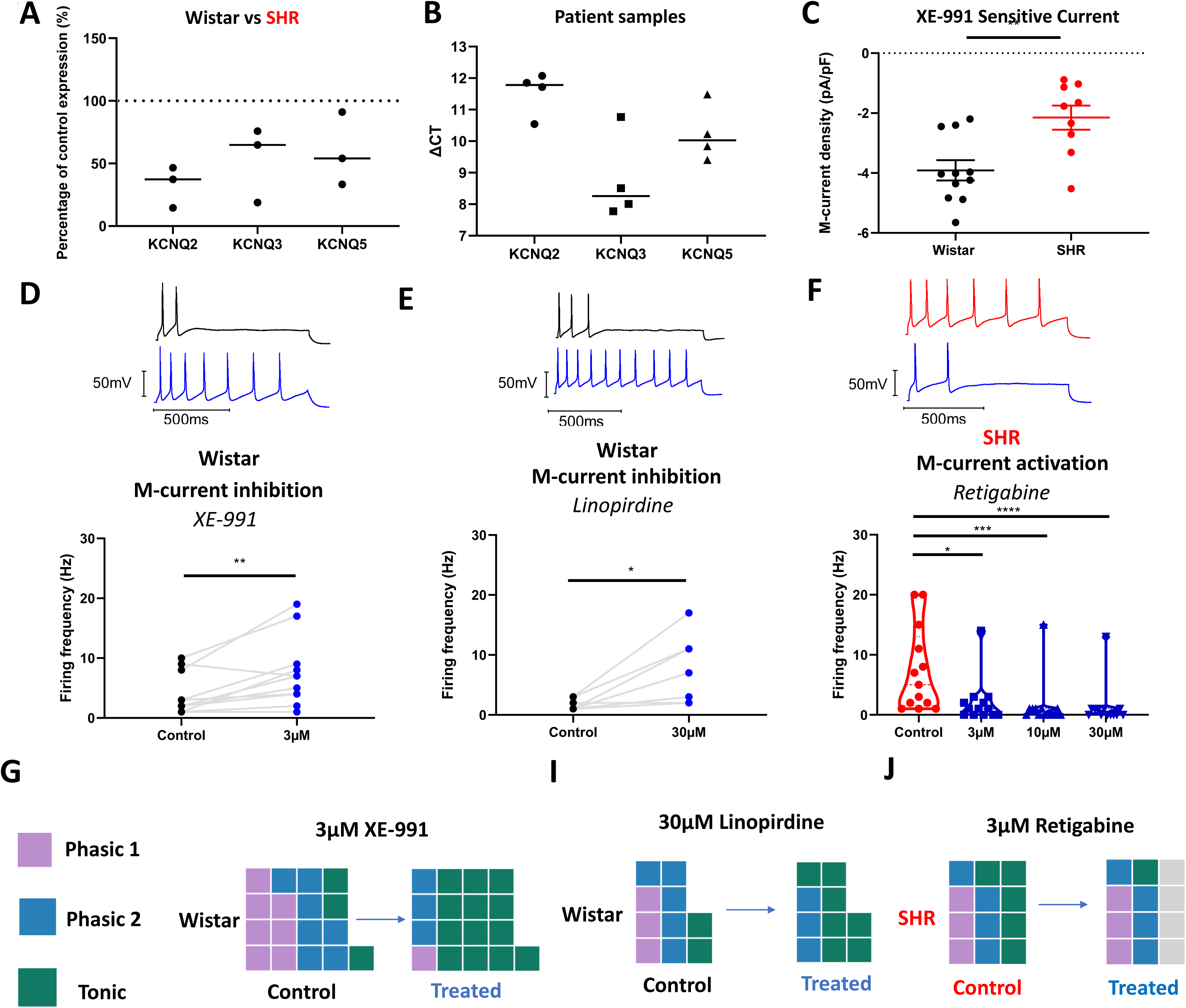
M-current is downregulated at a functional level in the stellate ganglia, and pharmacological manipulation of M-current can reverse or induce the firing rate phenotype observed in the SHR. (A) RT-qPCR reveals a downregulation of KCNQ2, KCNQ3 and KCNQ5 expression in the left stellate ganglia of 5-6 week old prehypertensive SHR (Median) (Percentage decrease; KCNQ2, 62.73%; KCNQ3, 35.16%; KCNQ5, 45.88%) (Wistar, n = 4; SHR, n = 3). (B) KCNQ2, KCNQ3 and KCNQ5 subunit expression was confirmed in patient samples of stellate ganglia, taken from donors (Median) (ΔCt; KCNQ2, 11.78; KCNQ3, 8.259; KCNQ5, 10.03; n = 4). (C) M-current density was revealed to be decreased in the SHR as determined by a decrease in XE-991 sensitive current, via a step protocol from −20 mV to −50 mV (Median ± IQR) (Wistar, −4.032 pA/pF, n = 11; SHR, −1.770 pA/pF, n = 9) (Mann-Whitney test, p = 0.0074). (D) M-current inhibition by 3 µM XE-991 significantly increased maximum firing rate induced by current injections in the range 10-200 pA (Median) (Control, 3 Hz; Treated, 7 Hz) (Wilcoxon Test, n = 12, p = 0.0059). (E) M-current inhibition by the alternate inhibitor 30 µM linopirdine also induced a significant increase in firing rate (Median) (Control, 1 Hz; Treated, 5 Hz) (Wilcoxon Test, n = 8, p = 0.016). (F) M-current activation in SHR neurons by retigabine significantly reduced maximum firing rate at all tested doses (Median) (Control, 5 Hz; 3 µM Retigabine, 1 Hz; 10 µM Retigabine, 0 Hz; 30 µM Retigabine, 0 Hz) (Friedman Test, n = 13, p<0.0001) (Dunn’s Multiple comparisons; Control vs 3 µM, p = 0.018; Control vs 10 µM, p = 0.0002; Control vs 30 µM, p < 0.0001). (G)(H) A graphical representation of subtype changes following the application of M-current inhibitors 3 µM XE-991 and 30 µM linopirdine in Wistar neurons. A trend towards a predominantly tonic firing subtype was observed in both cases. (I) A graphical representation of the effect of M-current activator Retigabine on SHR firing rate, which prevents firing in a third of tested neurons (Grey squares).

M-current (I_M_) was analysed through deactivation curves, in this case applied from a holding potential of −25 mV to −55 mV, allowing for a relaxation of current corresponding to I_M_. These recordings were made in perforated patch, as I_M_ is known to rundown in whole cell recordings. To confirm that the current measured was I_M_ we subtracted and measured current that was inhibited by I_M_ inhibitor 10 µM XE-991 (Wang *et al*., 2000). These data were then normalised to cell capacitance. By this measure, I_M_ was shown to be functionally present in stellate ganglia neurons and to be downregulated in SHR relative to Wistar (Figure 3C).

### M-current inhibition increased the excitability of Wistar stellate ganglia neurons

M-current pharmacology was used to assess its role in firing rate alongside other electrophysiological parameters in stellate ganglia neurons. M-current inhibition by 3 µM XE-991 (Wang *et al*., 2000) caused a significant increase in Wistar stellate ganglia neuron firing rate, measured as the maximum firing rate of tested neurons within a stimulation range of 0-200 pA (Figure 3D). When these data were visualised in terms of neuronal subtypes we observed an increase in tonic neurons after XE-991 application and a reduction in phasic 1 and phasic 2 subtypes (Figure 3K). This was supported by a significantly decreased rheobase (Supplementary figure 7D) and an accompanying depolarisation of resting membrane potential (Supplementary figure 7A). Similar results were seen after application of 30 µM Linopirdine, an alternative M-current inhibitor at a dose comparable in efficacy to 3 µM XE-991 (Costa and Brown, 1997). Linopirdine increased maximum firing rate in Wistar neurons (Figure 3E). This also caused a reduction in phasic 1 and phasic 2 subtypes, and an increase in tonic neurons (Figure 3H). As with XE-991, Linopirdine also reduced rheobase amplitude (Supplementary figure 7E) and depolarised the resting membrane potential of these neurons (Supplementary Figure 7B).

### M-current activation reduced excitability of SHR and Wistar stellate ganglia neurons

Since I_M_ was shown to be reduced, but not entirely absent via 10 µM XE-991-substracted deactivation curves (Figure 3C), we also tested whether increasing SHR M-current via the activator retigabine (Main *et al*., 2000) would be sufficient to reduce firing rate. We found that retigabine significantly reduced SHR stellate ganglia neuron maximum firing rate at all 3 doses tested (Figure 3F). These data were visualised for maximal firing rate as neuronal subtypes in Figure 3I, where the number of tonic neurons decreases, and 4 cells were prevented from firing at 3 µM retigabine in the stimulation range 0-200 pA. These observations were accompanied by a hyperpolarisation of the resting membrane potential of these neurons at all doses tested (Supplementary figure 7C). Retigabine increased rheobase amplitude at 3 µM retigabine in neurons for neurons that still fired within 0-200 pA stimulation (Supplementary Figure 7F), but at higher doses too few neurons still fired in this range to allow for a quantitative comparison of rheobase amplitude.

### I_Na_, SK channels and K_v_2.1 also modulate SHR stellate neuron firing rate

Using scRNAseq and a series of pharmacological inhibitors, we screened a range of calcium and voltage-gated ion channels with a known role in determining the firing rate of other neuronal populations. In Figure 4A the patterns of expression for key channels implicated in firing rate (Supplementary figure 2D) are shown against cell populations highlighted in Wistar and SHR stellate ganglia (Figure 2A; Figure 2B). These channel subunits were chosen based upon the expression of these subunits in stellate ganglia neurons, availability of selective pharmacology and a demonstrated role in firing rate in other neuronal populations. Other channels implicated in firing rate or other cell processes are highlighted in supplementary Figure 3. For the chosen channels, either 1 or 2 pharmacological inhibitors were applied to SHR neurons to observe any effect on firing rate (Figures 4-5; Table 2). Pharmacological inhibitors which had a significant effect on the firing rate of these neurons are shown in Figures 4-5 and non-significant inhibitors are shown in Table 2.

**Table 2.** Effect of other channel inhibitors on SHR neuron firing rate. All channels included are non-significant for all stated current injection amplitudes (50, 100, 150 pA or maximum firing rate in response to any 10 pA current step between 0-200 pA). All tested by Wilcoxon tests.

|  | HCN |  | T-type |  | Ca2+ free | BK | CaCC | Nav1.7 |  | KNa |
| --- | --- | --- | --- | --- | --- | --- | --- | --- | --- | --- |
| Drug | Ivabradine | ZD7288 | TTA-A2 | TTA-P2 | EGTA | Iberiotoxin | Ani9 | Protx-III | Huwentoxin-IV | Lithium |
| Concentration | 3μM | 1μM | 500nm | 1μM | 5mM | 100nM | 1μM | 50nM | 50nM | N/A |
| Perforating agent | Amphotericin | Amphotericin | Amphotericin | Amphotericin | Amphotericin | Amphotericin | Amphotericin | Amphotericin | Amphotericin | Whole cell |
| Δ Firing rate 150pA | -1 | 0 | 0 | -1 | -1 | 0 | 0.5 | 0 | 2.5 | 0 |
| *P value* | *0.67* | *0.5* | *>0.99* | *0.60* | *0.59* | *0.81* | *0.87* | *0.38* | *0.11* | *0.84* |
| Δ Firing rate 100pA | 0 | 0.5 | 0 | -0.5 | 1.000 | 0 | 0.5 | 0 | 0.5 | 0 |
| *P value* | *0.75* | *0.95* | *0.25* | *0.89* | *0.22* | *0.9414* | *0.5469* | *0.44* | *0.19* | *0.5* |
| Δ Firing rate 50pA | 0 | 0.5 | 1 | 0 | 0 | 0 | 0.5 | -0.5 | 0 | 0 |
| *P value* | *0.19* | *0.25* | *0.16* | *>0.99* | *0.094* | *>0.9999* | *0.5156* | *0.63* | *0.5* | *0.25* |
| Δ Maximum firing rate | 0 | 0 | 0 | -0.5 | 0.5 | 0 | 1 | -0.5 | 1 | 0 |
| *P Value* | *0.13* | *>0.99* | *0.63* | *0.51* | *0.61* | *0.60* | *0.29* | *0.67* | *0.19* | *0.81* |
| n | 10 | 8 | 9 | 10 | 12 | 12 | 10 | 8 | 8 | 10 |

**Figure 4.**
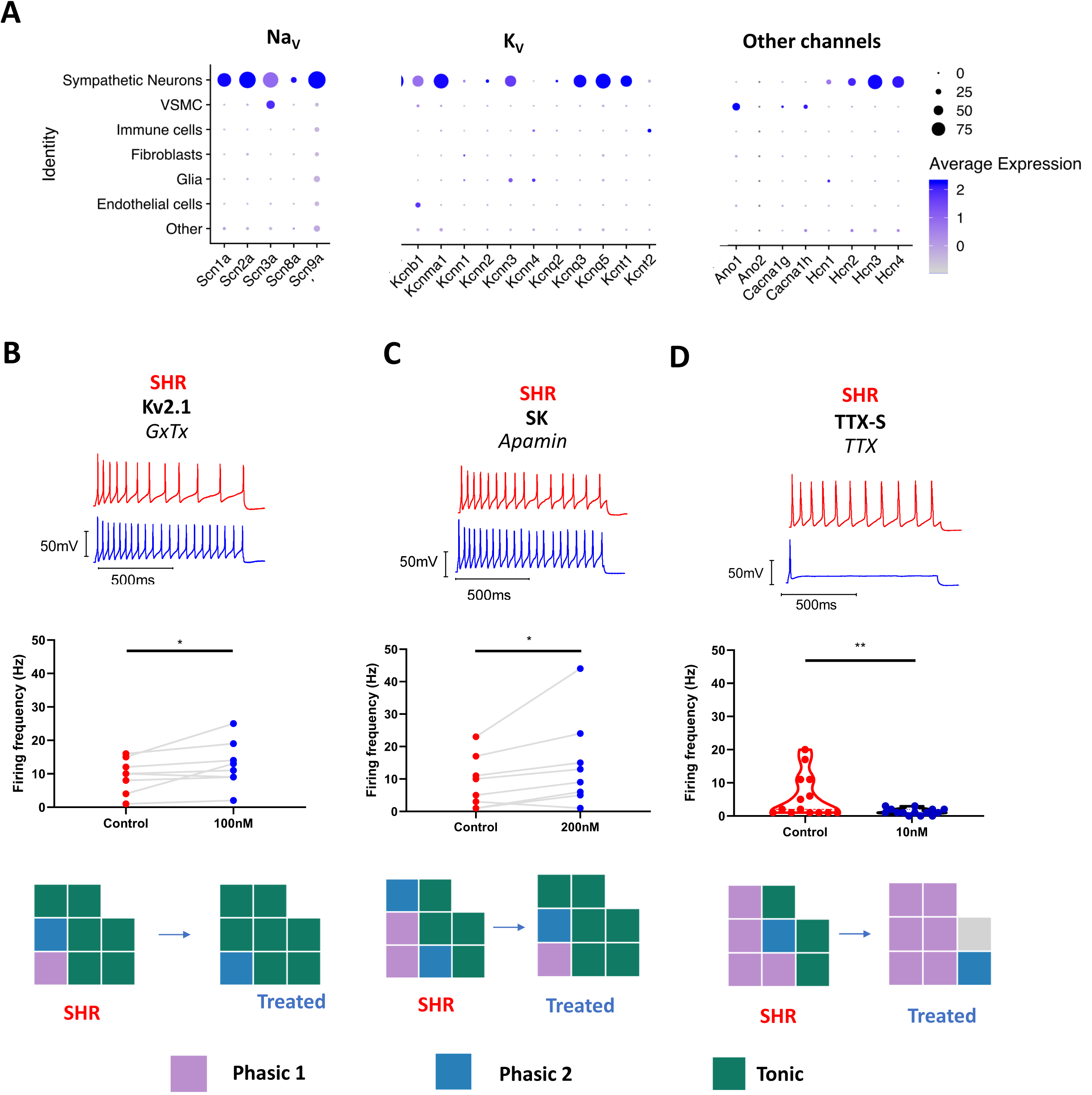
Single cell RNA-sequencing and a panel of pharmacological inhibitors were used to determine remaining channels involved in SHR enhanced firing, that may be targetable for the reduction of aberrant sympathetic hyperactivity or key to the SHR sympathetic pathology. (A) Single cell RNA-sequencing was used to identify the cell specific expression patterns of a range of channel subunits which are typically implicated in the control of firing rate in other neuronal populations. Here percentage expressed indicates the number of cells per cluster identity exhibiting transcript expression and average expression refers to the average expression per cluster identity. (B) Kv2.1 inhibition by 100 nM Guangxitoxin (GxTx) significantly increased maximum firing rate in SHR neurons (Median) (Control, 10 Hz; Treated, 12 Hz) (Wilcoxon test, n = 8, p = 0.039). (C) SK channel inhibition by 200 nM apamin caused a significant increase in firing rate (Median) (Control, 7.5 Hz; Treated, 11 Hz) (Wilcoxon test, n = 8, p = 0.016). (D) Generally targeting TTX-sensitive I_Na_ by 10 nM TTX caused a significant decrease in firing rate (Median) (Control, 2 Hz; 10 nM, 1Hz) (Wilcoxon test, n = 14, p = 0.0078).

**Figure 5.**
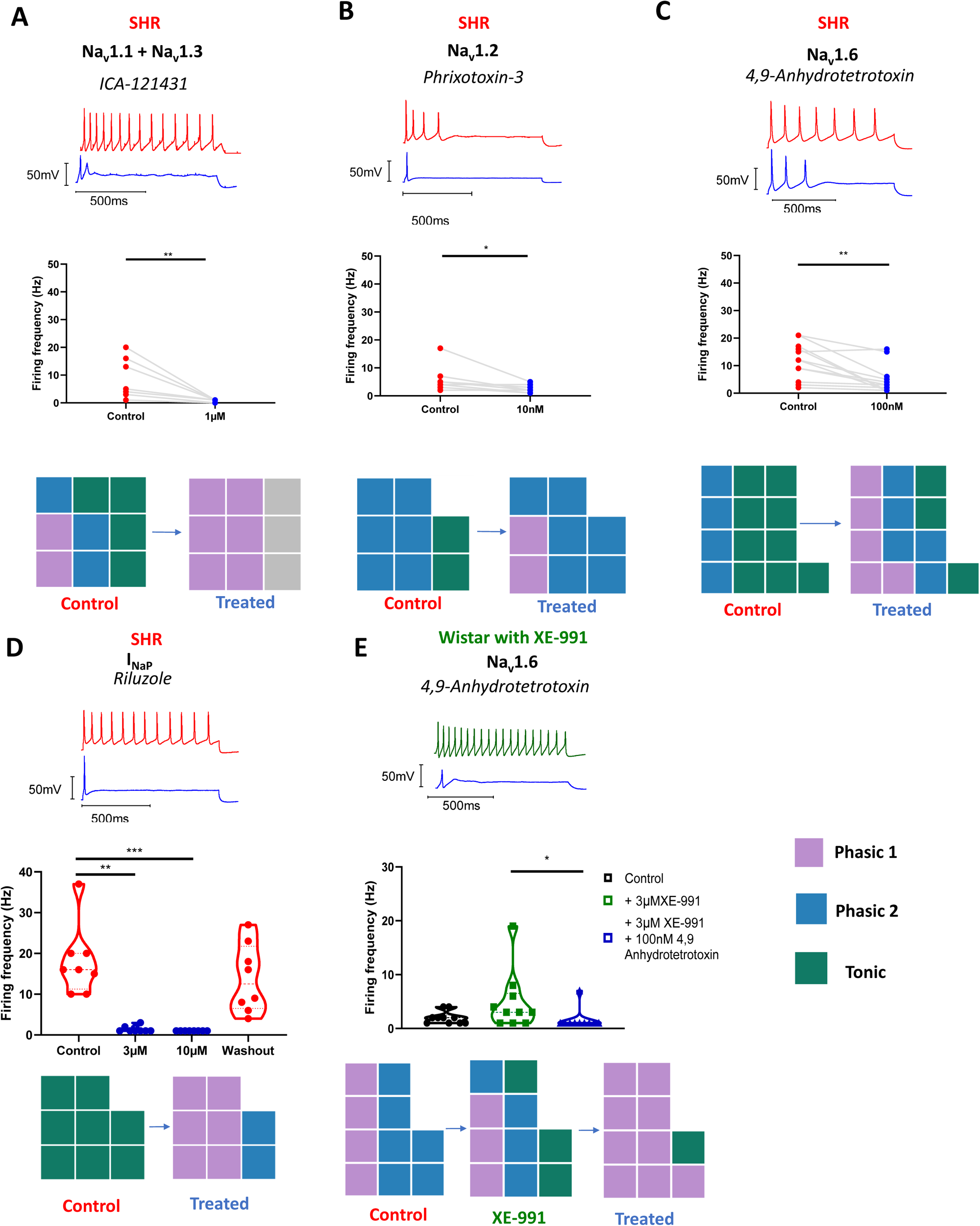
Exploring the role of Na_V_ subtypes in SHR firing rate. The effect of specific Na_V_ subtype and conductance state inhibitors were investigated in SHR stellate ganglia neurons. (A) Na_v_1.1 inhibition by 1 µM ICA-121431 in SHR neurons significantly reduced maximum firing rate in SHR neurons (Median) (Control, 4 Hz; Treated, 1 Hz) (Wilcoxon test, n = 9, p = 0.0039). (B) Na_v_1.2 inhibition by 10 nM Phrixotoxin-3 in SHR neurons significantly reduced maximum firing rate in SHR neurons (Median) (Control, 4.5 Hz; Treated, 2 Hz) (Wilcoxon test, n = 8, p = 0.0313). (C) Nav1.6 inhibition by 100 nM 4,9-Anhydrotetrodotoxin significantly reduced maximum firing rate in SHR neurons (Median) (Control, 12 Hz; Treated, 3 Hz) (Wilcoxon test, n = 13, p = 0.0015). (D) I_NaP_ inhibition by 3-10 µM Riluzole in SHR neurons reduced firing rate (Median) (Control, 16 Hz; 3 µM, 1 Hz; 10 µM, 1 Hz; Washout, 12.5 Hz) (Friedman test, n = 8, p < 0.0001) (Dunn’s multiple comparisons test; Control vs 3 µM, p = 0.003; Control vs 10 µM, p = 0.007). (E) Nav1.6 inhibition significantly reduced firing rate in M-current (3 µM XE-991) inhibited Wistar neurons (Median) (Control, 2 Hz; XE-991 treated, 3 Hz; XE-991 and 4,9-Anhydrotetrotoxin treated, 1 Hz) (Friedman test, n = 10, p = 0.0084) (Dunn’s Multiple comparisons test; XE-991 treated vs XE-991 + 4,9-Anhydrotetrotoxin, p = 0.038).

Low dose (10 nM) Tetrodotoxin (TTX) (Tucker *et al*., 2012) significantly reduced firing rate in SHR neurons (Figure 4D). In a separate population of SHR neurons, high dose TTX (300 nM) was shown to prevent firing in all tested SHR neurons (Supplementary figure 8F), confirming the absence of a large TTX-insensitive sodium current (I_Na_).

Two compounds tested, K_v_2.1-2.2 inhibitor, Guangxitoxin (Li *et al*., 2013) and SK channel inhibitor, Apamin (Kawai and Watanabe, 1986), increased SHR stellate ganglia neuron firing rate (Figures 4B-C). 100 nM Guangxitoxin (Figure 4B) significantly increased SHR firing rate from 10 Hz at baseline to 12 Hz, whereas 200 nM Apamin (Figure 4C) increased median firing rate from 7.5 Hz to 11 Hz. The effect of these inhibitors on firing rate subtypes is shown in Figures 4B-4C, where we observed an increase in the number of tonic firing neurons after 100 nM Guangxitoxin or 200 nM Apamin addition (Figure 4B-C).

### Specific Na_v_ subunit inhibitors reduce SHR stellate neuron firing rate

Using a panel of selective inhibitors, we inhibited Na_V_ subunits shown to be present in the stellate ganglia via scRNAseq (Figure 4A). The Na_v_1.1 and Na_v_1.3 inhibitor, ICA-121431 (McCormack *et al*., 2013), was tested for its effect on SHR stellate ganglia neuron firing rate. 1 µM ICA-121431 significantly reduced SHR firing rate (Figure 5A), and converted a majority of neurons to phasic 1 (Figure 5A). The Na_v_1.2-1.3 and Na_v_1.5 inhibitor, phrixotoxin-3 is specific for Na_v_1.2 at the tested dose, 10 nM, and was therefore utilised as a selective Na_v_1.2 inhibitor (Bosmans *et al*., 2006). 10 nM phrixotoxin-3 significantly reduced SHR neuron firing rate (Figure 5B) and prevented tonic firing (Figure 5B). Inhibition of Nav1.6 by 4,9-Anhydrotetrodotoxin (Rosker *et al*., 2007) significantly reduced firing of SHR neurons (Figure 5C). We also tested 100 nM 4,9-Anhydrotetrotoxin (Rosker *et al*., 2007) on XE-991 inhibited Wistar neurons, and found that, similar to in the SHR neurons, it significantly decreased firing rate (Figure 5E) and converted the majority of the tested neurons to phasic 1 (Figure 5D). The persistent Na_v_ channel inhibitor riluzole (Urbani and Belluzzi, 2000) inhibited firing in the tested population and reduced most neurons to phasic 1 at the lowest dose tested, 3 µM (Figure 5D).

Membrane depolarisation can limit Na_v_ availability through increasing Na_V_ inactivation (Ulbricht, 2005). To ensure the depolarised resting membrane potential observed in the SHR (Figure 1D) did not limit SHR firing rate, we applied a series of negative 1 second current injections in the range −10 to −100 pA followed immediately by a stimulatory 1 second 150 pA current injection. By this method, we found that there was no significant difference between these current steps or a 0 pA control prepulse in SHR neurons (Supplementary Figure 8G).

Further to these data, we observed that Na_v_1.8 inhibitors, which have previously been shown to inhibit stellate ganglia function *in vivo* (Yu *et al*., 2017), are unlikely to act through this mechanism. 100 nM A803467 (Jarvis *et al*., 2007) and 300 nM A887826 (Zhang *et al*., 2010) inhibited firing at tested doses (Supplementary Figure 8A) with no observed change in membrane potential (Supplementary Figure 9C-D). However, a third Na_v_1.8 inhibitor PF04885614 (1 µM) did not change firing rate (Supplementary figure 9B) or resting membrane potential (Supplementary figure 9E). We also found that Na_v_1.8 was not present by RT-qPCR nor scRNAseq, an observation supported by previous work in the superior cervical ganglion (SCG) and the lack of TTX insensitive Na_v_ (Supplementary Figure 8F).

### Tonic neurons have less I_M_ and more I_Na_

By studying action potential kinetics and I_M_ density between subtype populations, we aimed to gain further insight into the mechanisms behind these subtypes. First, we found that in perforated patch clamp recordings input resistance was higher in tonic neurons than phasic 1 and phasic 2 (Figure 6A). Indirect measures of I_Na_, action potential upstroke and action potential amplitude, taken from whole cell recordings of single action potentials induced by a threshold 10 ms current injection were used to highlight any differences between subtypes. Whole cell recordings were used to reduce the effect of higher R_s_ values encountered with perforated patch on action potential kinetics. These data reveal higher I_Na_ in tonic and phasic 2 populations than in phasic 1 as measured by amplitude (Figure 6B) or upstroke (Figure 6C).

**Figure 6.**
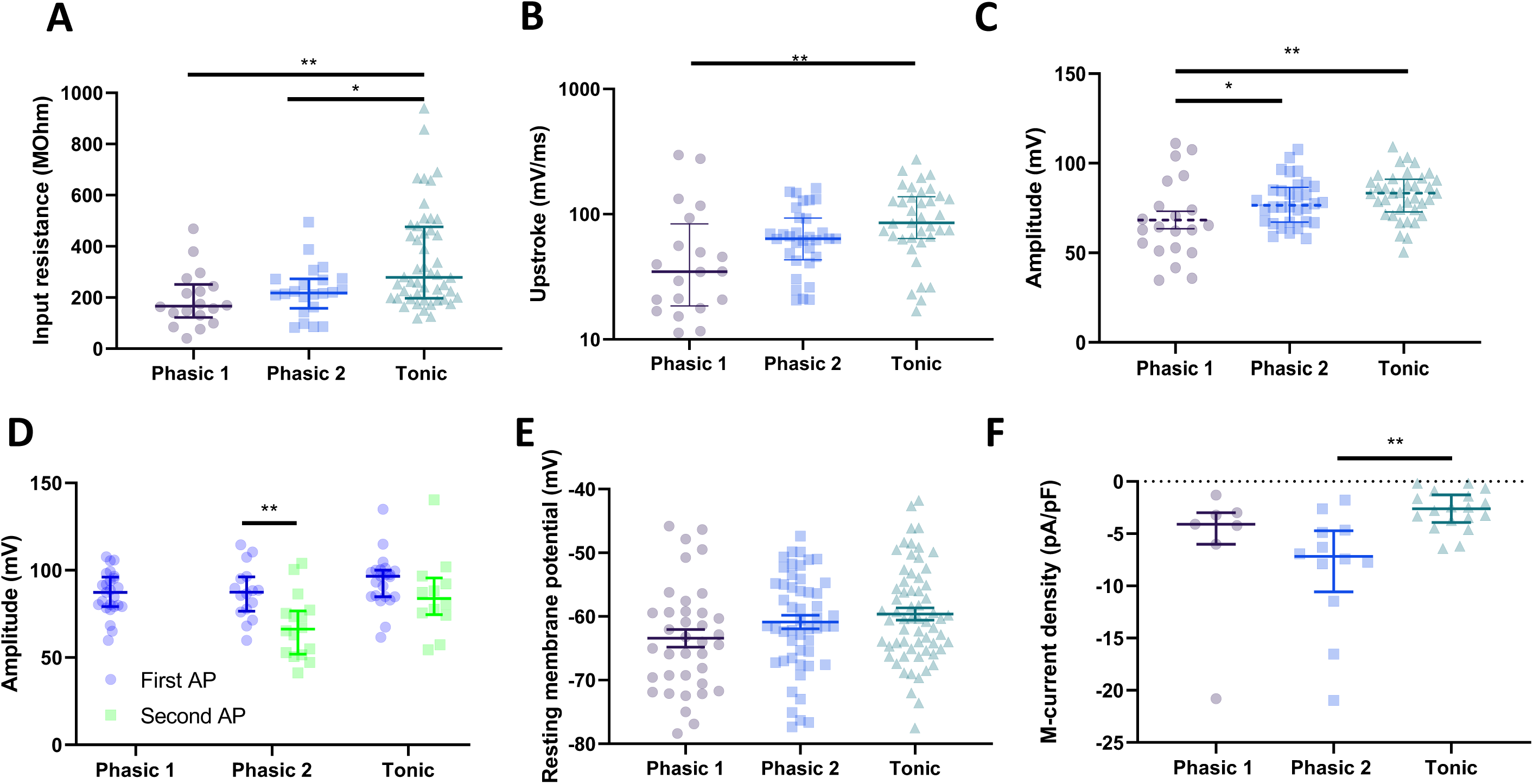
Neural subtypes correlate with measures of I_M_ and I_Na_. (A) Input resistance was significantly higher in tonic neurons than in phasic 2 and phasic 1 neurons (Median ± IQR) (Phasic 1, 166.5 MΩ, n = 18; Phasic 2, 218.2 MΩ, n = 22; Tonic, 279 MΩ, n = 48) (Kruskal-Wallis test; p = 0.0006) (Dunn’s multiple comparisons test; Phasic 1 vs Tonic, p = 0.011; Phasic 2 vs Tonic, p = 0.046). (B) Upstroke velocity for phasic 1, phasic 2 and tonic neurons revealed a higher Tonic velocity (Median ± IQR) (Phasic 1, 34.71 mV/ms, n = 20; Phasic 2, 63.65 mV/ms, n = 32; Tonic, 85.38 mV/ms, n = 37) (Kruskal-Wallis test; p = 0.0018) (Dunn’s multiple comparisons test; Phasic 1 vs Tonic, p = 0.0012). (C) Action potential amplitude was significantly higher for phasic 2 and tonic neurons than in phasic 1 neurons (Mean ± SEM) (Phasic 1, 68.30 ± 4.49 mV, n = 21; Phasic 2, 78.25 ± 2.25 mV, n = 32; Tonic, 81.96 ± 2.14mV, n = 37) (One-way ANOVA; p = 0.0071) (Holm-sidak’s multiple comparisons test; Phasic 1 vs Phasic 2, p = 0.049; Phasic 1 vs Tonic, p = 0.0054). (D) The amplitude of the second action potential in a train was lower in phasic 2 neurons, but not tonic suggesting an inability to support repetitive firing in phasic 2 neurons (Mean ± SEM) (Phasic 1, First AP 86.67 ± 2.48 mV, n = 25; Phasic 2, First AP 87.34 ± 3.8 mV, Second AP 67.76 ± 4.58 mV, n = 16; Tonic, First AP 93.67 ± 3.22 mV, Second AP 85.52 ± 6.48mV, n = 22) (Multiple t-test’s; Phasic 2, p = 0.0026; Tonic, p = 0.22). (E) Resting membrane potential for the three electrophysiological subtypes (Mean ± SEM) (Phasic 1, −63.43 ± 1.38 mV, n = 37 ; Phasic 2, −60.88 ± 1.07 mV, n = 51; Tonic, − 59.62 ± 0.96mV, n = 63) (One-way ANOVA, p = 0.065). (F) M-current density was lower in Tonic firing neurons than phasic 2 neurons (Median±IQR) (Phasic 1, −4.09 pA/pF, n=7; Phasic 2, −7.18 pA/pF, n=12; Tonic, −2.62 pA/pF, n=18) (Kruskal-Wallis test; p = 0.0019) (Dunn’s multiple comparisons test; Phasic 2 vs Tonic, p = 0.0012).

Further to these data, by comparing the amplitude of the first and second action potential elicited by a threshold 10 pA 1000 ms current injection, we found that in phasic 2, but not tonic neurons, there was a significant decrease in the second action potential amplitude (Figure 6D). We confirmed that these parameters were I_Na_ dependent via the effect of low dose TTX on single action potentials as measured by action potential upstroke in perforated patch (Supplementary Figure 7A) or amplitude (Supplementary Figure 7B). There were no significant differences between strains overall in action potential upstroke (Supplementary Figure 7C) or amplitude (Supplementary Figure 7D) measured by whole cell patch clamp recordings.

There were no significant differences in resting membrane potential between the three subtype groups when compared via perforated patch (Figure 6E). However, there was a trend towards a more depolarised resting membrane potential in phasic 2 and tonic neurons. When viewed per electrophysiological subtype, we observed significantly less I_M_, as determined by deactivation curves, in tonic firing neurons than phasic 2 neurons (Figure 6F).

## Discussion

We report four primary novel findings. First, sympathetic stellate ganglia neurons from the prehypertensive SHR are hyperexcitable, which manifests as a higher firing rate, depolarised resting membrane potential and reduced rheobase. Secondly, I_M_ is downregulated in the stellate ganglia neurons of the SHR and this is the causative mechanism for membrane hyperexcitability. Thirdly, M-current is conserved in human stellate ganglia. Fourthly, hyperexcitability can be curbed either by elevation of remaining I_M_ or via reduction of I_Na_, via non-selective or selective inhibition of Na_v_1.1-1.3, Na_v_1.6 or I_NaP_. These targets may provide therapeutic utility for reducing cardiac sympathetic neuron activity pharmacologically without resorting to surgery.

Sympathetic hyperactivity is well documented in hypertension, myocardial infarction and heart failure and contributes to the pathogenesis of disease progression (Herring, Kalla and Paterson, 2019). The phenotype we observed of increased neuronal firing rate is also seen alongside increased cardiac noradrenaline spill over in human hypertension (Esler, 2000), and we suggest that this is at least in part secondary to the intrinsic properties of post-ganglionic stellate neurons rather than solely being driven by higher centres. Previous work in the SHR model has reported repetitive firing in neurons of the superior cervical ganglia (SCG), although there was no mechanistic data underlying this observation (Robertson and Schofield, 1999). A following study suggested that enhanced firing may be due to changes in I_M,_ although again there was limited biophysical evidence for the speculation(Yarowsky and Weinreich, 1985). Subsequent work suggested the altered phenotype is due to changes in the A-type K^+^ current (I_A_), which was reported to be larger in SCG neurons of the SHR (Robertson and Schofield, 1999), even though I_A_ is similar in phasic and tonic neurons (Wang and McKinnon, 1995). Moreover, we observed no changes in I_A_ encoding transcript expression by scRNAseq (Figure 2D; Supplementary Figure 2D) between SHR and Wistar stellate neurons. Notwithstanding the fact that a plethora of potassium channels have been documented in other ganglia (Dixon and McKinnon, 1996), we felt it was best to use an unbiased scRNAseq approach to identify molecular targets.

### Role of I_M_

The increased firing rate and neuronal subtypes we observe in the SHR (Figure 1) fits well with the known characteristics of I_M_ (Brown and Adams, 1980). I_M_ is a non-inactivating inhibitory K^+^ current, and provides a restriction upon firing, but only after its considerable activation period(Brown and Adams, 1980). This fits with both a general increase in firing rate (Figure 1A; Figure 1B), and a time-dependent phenotype (Figure 1C). I_M_ has a powerful effect on neuronal resting membrane potential, as demonstrated in Supplementary Figure 5A-C, therefore the loss of I_M_ in the SHR is likely to cause the depolarisation of the resting membrane potential in the SHR (Figure 1D). Notably, reversing this depolarisation alone does not appear to alter firing rate (Supplementary figure 8F). I_M_ inhibition also reduces rheobase (Supplementary Figure 7D-E), which again fits with the phenotype observed (Figure 1E).

Importantly, we demonstrate that inhibition of I_M_ was enough to recapitulate the SHR phenotype in the normal rat. Prior work has investigated the systemic effect of I_M_ modulators *in vivo* and highlighted a role for autonomic I_M_ at a systems level in Wistar controls and SHR (Berg, 2016, 2018). This would be expected based upon our observation of I_M_ having a dominant role in stellate ganglia neuron function *in vitro* (Figure 3) and our transcriptional and functional data highlighting I_M_ as a powerful driver of hyperactivity in the SHR. Retigabine has been demonstrated to inhibit noradrenaline release and reduce neurotransmission from SCG neurons, (Hernandez *et al*., 2008). Finally, human genome wide association studies have found two M-current single nucleotide polymorphisms in KCNQ3 (rs138693040-T) (Méndez-Giráldez *et al*., 2017) and KCNQ5 (rs12195276-T) (Evangelou *et al*., 2018) to be significantly associated with variation in the electrocardiographic QT interval and pulse pressure respectively, both variables which are modulated by the sympathetic nervous system^2^.

### Role of I_Na_

As observed for other neuron populations (Bennett *et al*., 2019), I_Na_ modulation also appears to be a powerful regulator of firing rate in SHR stellate ganglia neurons (Figure 4D; Figure 5). The pattern of expression for Na_v_ subunits is different than for sensory or central neurons, with a stellate ganglia neuron expressing Na_v_1.1-1.3 and Na_v_1.6-1.7 (Figure 4A). Stellate ganglia neurons express peripheral channel Na_v_1.6, but lack Na_v_1.8 and Na_v_1.9 for which subpopulations of sensory peripheral neurons are notable (Bennett *et al*., 2019). Inhibition of all Na_v_ subunits identified by scRNAseq (Figure 4A), apart from Na_v_1.7, significantly reduced SHR neuron firing (Figure 6). This is useful clinically in terms of finding a tolerable inhibitor as a range of Na_v_ subtypes could be targeted to reduce I_Na_.

Of interest, there are case reports of thoracic epidural anaesthesia (using Na^+^ channel blockers) being successfully used in patients experiencing recurrent, life threatening ventricular tachycardias to block sympathetically driven arrhythmia as a bridge to surgical cardiac sympathetic denervation via stellectomy (Meng *et al*., 2017; Herring, Kalla and Paterson, 2019). Na_v_1.7 inhibition does not affect firing rate by either tested inhibitor (Table 2), suggesting that Na_v_1.7 does not contribute to firing rate in the SHR. Na_v_1.7 undergoes sustained inactivation, unlike Na_v_1.6 for example (Bennett *et al*., 2019), and is therefore less likely to support sustained firing. Na_v_1.7 inhibitors have been billed as a treatment for neuropathic pain, and have seen much academic and industrial attention. Thus the lack of effect on cardiac sympathetic firing can be regarded as a positive indicator that these compounds are unlikely to have sympathetic side effects.

Na_v_1.1 (*Scn1a*), Na_v_1.2 (*Scn2a*) and Na_v_1.7 (*Scn9a*) mRNA expression is downregulated in the SHR (Table 1), a possible compensatory mechanism, which does not appear to negatively affect SHR neuron firing rate (Figure 1), action potential amplitude (Supplementary Figure 8D) or action potential upstroke (Supplementary Figure 8C). Therefore, these changes either do not cause a reduction in channel protein or that the reduction in I_Na_ is minimal.

A803647, an inhibitor of the TTX-resistant Na_v_ subunit, Na_v_1.8, has been previously shown to have a powerful inhibitory effect on arrhythmogenesis following stellate ganglia stimulation when applied locally *in vivo* (Yu *et al*., 2017). This was originally attributed to an effect on stellate ganglia neuron Na_v_1.8 function, however we find no molecular evidence of its expression (Figure 4A) nor could firing occur after 300nM TTX (Supplementary figure 8F). Our data show that the effect of this inhibitor is likely to be through an off-target reduction of I_Na_, as shown on a cellular level for A803647 and A887826 (Figure 4A; Supplementary Figure 9) and suggested by other *in-vivo* studies (Stone *et al*., 2013).

### Role of other ion channels

We found no expression differences for calcium-activated channels encoding genes in our scRNAseq dataset (Figure 2D). Further to this, removal of extracellular calcium had no net effect on SHR neuron firing rate (Table 2). Some calcium-activated channels may have a role in fine tuning stellate neuron firing rates, as shown by the increased firing rate after Apamin inhibition of SK channels. Apamin has been reported to increase firing rate in SCG neurons, but to a lesser extent than observed here (Figure 4D) (Kawai and Watanabe, 1986). In the SCG, BK channels may play a key role in determining the spiking response to bradykinin and sensitivity to NGF (Vivas, Kruse and Hille, 2014). Our scRNAseq data supports BK channel expression (Figure 4A) and we observed an increase in action potential width similar to that previously described in dopaminergic neurons (Kimm, Khaliq and Bean, 2015). We therefore suggest that BK channels are likely to have a role in stellate ganglia neurons independent of firing rate. The only other conductance we found to be expressed in stellate ganglia neurons which influenced firing rate was K_v_2.1, which increased firing rate, as has been reported in the SCG ^11^.

A range of neuronal subtypes based on firing properties have been reported including phasic 1, phasic 2 and tonic firing neurons (Wang and McKinnon, 1995), although the electrophysiological basis for these phenotypes is unclear. Whilst we observe the presence of all 3 subtypes in the stellate ganglia (including ∼20% tonic neurons), the SCG contains only phasic 1 and phasic 2, whilst tonic neurons have been observed in the coeliac (∼58%) and superior mesenteric (∼85%) ganglia (Wang and McKinnon, 1995). We also observed that neurons of the Wistar stellate ganglia have a much larger input resistance (218.2 MΩ) in comparison to the majority of measurements from SCG (Mouse 109.7 MΩ, 75-85 MΩ; Rat 389.9 MΩ), thoracic (Mouse 117.9 MΩ) and coeliac (Mouse 161.9 MΩ; Guinea pig 116.9 MΩ) ganglia (Jobling and Gibbins, 1999; Anderson, Jobling and Gibbins, 2001; Lamas, Reboreda and Codesido, 2002; Martinez-Pinna *et al*., 2018). Previous studies have also observed more depolarised resting membrane potentials in other sympathetic ganglia, with our estimated value in Wistar of −68.04 mV being more hyperpolarised than most recordings from other ganglia including the SCG (Mouse SCG −49.9 mV, −54 mV; Rat SCG −58.3 mV) (Jobling and Gibbins, 1999; Martínez-Pinna, McLachlan and Gallego, 2000; Lamas, Reboreda and Codesido, 2002). However, some of these differences may be attributed to interspecies as well as inter-ganglia differences.

### I_Na_ and M-current in electrophysiological subpopulations

Our pharmacological data is consistent with the hypothesis that the behaviour of stellate ganglia phasic 1 neurons is due to relatively low I_Na_ compared to phasic 2 and tonic neurons, which would limit sustained firing (Figure 5; Figure 7). Also, the difference between phasic 2 and tonic neurons is likely due to lower I_M_ with a spectrum of current amplitudes and associated firing rates. Others have also suggested a relationship between reduced I_M_ (Wang and McKinnon, 1995; Jia *et al*., 2008; Luther and Birren, 2009) and higher I_Na_ (Luther and Birren, 2009) in different neuronal populations, although we specifically define their role between the 3 sub-types and propose a link to pathology. The relative ease of pharmacologically converting these neural subtypes (Figures 3-5), suggests they are fluid and dynamic, and do not correspond to hardwired neuronal subtypes. This is supported by our scRNAseq data, where two main populations of sympathetic neurons appear (Figure 2B), which can be divided by one population expressing high levels of the co-transmitter neuropeptide-Y. Both groups express similar levels ion channels involved in determining firing rate (Supplementary figure 6) and do not correlate with the electrophysiological sub-types described. These transcriptome defined groups merit further study, but a detailed comparison is beyond the scope of this study.

Whilst downregulation of I_M_ provides an explanation for membrane hyperactivity, we have not identified a pathway by which this may occur. One possible explanation is that this results from continual presynaptic input, which contributes to ganglionic LTP (Cifuentes, Arias and Morales, 2013). It is also possible that other channels may be contributing to this phenotypic difference, but as the measured variables all correlate well with our observations of I_M_ inhibition in Wistar neurons, it seems highly likely that I_M_ downregulation alone is a major cause.

### Limitations

This study has several limitations. For example, the SHR model of hypertension has a genetic basis as well as being rodent physiology, which differs in several respects from man (Hasenfuss, 1998).To address the translatability of our work, we have confirmed that I_M_ subunits are expressed in human stellate ganglia indicating conservation of transcript (Figure 3B), but we have not been able to assess functionality. Indeed, RNA sequencing does not establish whether transcripts encode proteins that confer physiological function. Nevertheless scRNAseq was the best approach for this study, as bulk sequencing would also incorporate contaminating cell types, for example vascular cells which are known to have ion channel expression changes in the SHR (Jepps *et al*., 2011). It should be noted that we observed downregulation of all three M-current encoding subunits via RT-qPCR, but only downregulation of KCNQ5 by scRNAseq. For KCNQ2, the detected expression level is relatively low via scRNAseq (Figure 2C). For both KCNQ2 and KCNQ3 it is possible that downregulation of contaminating vascular SHR KCNQ2 and KCNQ3 is detected by our RT-qPCR of the whole stellate ganglion. Regardless, these data support a downregulation of I_M_ in stellate ganglia neurons of the SHR, with a functional reduction of I_M_ supporting this.

In conclusion, we have described in detail a phenotype of sympathetic hyperactivity in stellate ganglia neurons of the SHR and provided an electrophysiological framework for this observation, guided by scRNAseq of the stellate ganglia and human validation of key transcripts. Targeting certain ion channels in the stellate ganglia, such as M-current, may provide a reversible therapeutic opportunity to treat cardiac sympathetic hyper-responsiveness over and above interventions like surgical stellectomy.

## Methods

### Animal work declaration

Animal use complied with the University of Oxford local ethical guidelines and was in accordance with the Guide for the Care and Use of Laboratory Animals published by the US National Institutes of Health (NIH Publication No. 85-23, revised 2011) and the Animals (Scientific Procedures) Act 1986 (UK). Experiments were performed under British Home Office Project License PPL 30/3131 and P707EB251. All animals were ordered from Envigo and housed on a 12-hour day-night cycle. Animals were sacrificed via an overdose of pentobarbitone and confirmed via exsanguination according to schedule one of the Animals (Scientific Procedures) Act 1986 (UK).

### Animals

Male normotensive Wistar rats and prehypertensive SHR’s were culled at 5-6 weeks of age, at which age SHR’s possess a phenotype unaffected by prolonged hypertension as found in older animals. Moreover, around this age post-natal ion channel expression stabilizes in sympathetic ganglia (Hadley *et al*., 2003).

### Human stellate ganglia tissue

Stellate ganglia were collected from organ donors at the time of organ procurement as approved by the UCLA IRB: 12-000701. Written informed consent was provided by the patient or appropriate designee. This study complies with the Declaration of Helsinki.

### Cell isolation and culturing

Stellate ganglia were dissected and immediately transferred to ice-cold HEPES buffered L15 media (L1518, Thermofisher, US). Ganglia were then cut into 2 mm sections using surgical scissors, and enzymatically digested at 37 °C first using 1 mg/ml Collagenase IV (Worthington, US) in L15 for 25 minutes, followed by 30 minutes in 2 mg/ml Trypsin (Worthington, US) in Ca^2+^ and Mg^2+^ free Hanks buffered salt solution (Thermofisher, US). Enzymes were then inhibited using two washes of a blocking solution containing 10% FBS. The tissue was then suspended in a plating media containing Neurobasal plus media, B27 plus, 100 ng/ml 2.5s NGF, 25 mM glutamax and 50 units/ml Pen-strep and mechanically disrupted using a fire-blown glass pipette. The cell suspension was then plated onto Poly-D-lysine coated Fluorodish 35 mm dishes (WPI, US), which had been previously incubated for 2 hours with 1 ug/ml laminin, a concentration chosen to allow cell adhesion and survival, but limiting neurite outgrowth (Buettner and Pittman, 1991). The cells were then incubated at 37 °C with 5% CO_2_ for a period of 1-5 days in vitro before use. All datasets were recorded from at least 2 cultures, with each culture requiring 4 animals. Phase contrast microscopy was used to enable neuronal identification. Neurons were identified in dissociated culture based on their large size and circular somata relative to the surrounding cell types.

### Electrophysiological Data acquisition

All electrophysiological data were acquired using Winwcp (Version 5.4.0) and recorded via a Multiclamp 700B amplifier (Molecular Devices, US) with an axon digidata 1550A (Molecular devices, US) digitizer. All current clamp recordings were sampled at 10 kHz. M-current deactivation curves were sampled at 10 KHz.

### Perfusion

Cells were constantly perfused at a rate of 5-6 ml/min, drugs were applied via this perfusion system. Most drugs were continuously applied for a duration of 5 minutes before data acquisition, XE-991, substituted LiCl and Linopirdine were perfused for 10 minutes before recording. Recordings were performed at room temperature.

### Perforated patch-clamp recordings

Amphotericin B (0.48 mg/ml) was used as the perforating agent (Rae *et al*., 1991), recordings were initiated after low series-resistance electrical access was achieved (<30 MΩ) and stable for a period of five minutes. All voltage clamp recordings used 70% series resistance compensation and capacitance cancellation. Current clamp recordings were bridge balanced. All recordings were monitored throughout, and recordings with R_S_ changes > 20% were discarded. The external recording solution for both current clamp and voltage clamp recordings was as follows: 5.2 mM KCl, 140 mM NaCl, 1 mM MgCl_2_, 1.8 mM CaCl_2_, 10 mM HEPES, 10 mM D-Glucose. External solution pH was adjusted to 7.4 with NaOH. For removal of external calcium, calcium was excluded from this solution and 5 mM EGTA was added. The internal solution was composed as follows: 145 mM K^+^-Aspartate, 2.2 mM EGTA, 10 mM HEPES, 1 mM MgCl_2_. Internal pH was adjusted to 7.3 using KOH.

### Whole cell patch-clamp recordings

Whole cell action potential recordings with Rs values >12 MΩ discarded. Current clamp recordings were bridge balanced and membrane potentials were corrected for liquid junction potentials. All recordings were monitored throughout, and recordings with R_S_ changes > 20% were discarded. Single action potentials were evoked via a 10ms positive current injection, at the minimal required injection size. External solution composition was the same as for perforated patch. For LiCl substitution experiments, 140 mM LiCl was substituted for NaCl. The internal solution was as follows: 130 mM K^+^-Gluconate, 10 mM KCl, 10 mM HEPES, 10 mM Na+-Phosphocreatine, 4 mM MgATP, 0.3 mM Na_2_GTP. Internal pH was adjusted to 7.3 with KOH. To study K_Na_, NaCl was substituted for LiCl to allow an inward g_Li_ via Na_V_ channels but prevent activation of slick and slack channels by intracellular sodium (Kaczmarek, 2013). For Ca^2+^ free experiments, Ca^2+^ was removed from the external solution and 5 mM EGTA, a Ca^2+^ buffer, was added.

### qRT-PCR

Total RNA from whole flash frozen stellate ganglia was isolated using a RNeasy minikit (Qiagen, US) and immediately stored on dry ice before cDNA library preparation. For cDNA synthesis, Superscript IV VILO with ezDNase genomic DNA depletion (Thermofisher, US) was used, cDNA was then stored at −80°C until required. Taqman PCR primers were used for the transcript identification of KCNQ2 (Rn00591249_m1), KCNQ3 (Rn00580995_m1), KCNQ5 (Rn01512013_m1), SCN10A (Rn00568393_m1), where either GAPDH (Rn01775763_g1) or B2M (Rn00560865_m1) were used to normalize values via the ΔΔCT method (Livak and Schmittgen, 2001). Samples were measured on an ABI Prism 7000 (Thermofisher, US) as per the standard protocol for taqman.

### Cryosectioning and Immunohistochemistry

Freshly isolated stellate ganglia were immediately transferred to 4% paraformaldehyde for 1-2 hours, after which the tissue was incubated overnight in 20% sucrose-PBS at 4 °C, before embedding in OCT compound (Tissue-Tek). Tissue was then frozen and stored at −80 °C until cryosectioning the tissue as 12 µm sections. Slides were then permeabilized in 0.3% triton-X for 30 minutes at room temperature, before blocking for 2 hours in 1% BSA, 5% donkey serum. Sections were then incubated for 24 hours with primary antibodies at 4°C, followed by five 5 minute washes in PBS and 2 hours incubation with the relevant secondary antibodies. Sections were subsequently washed 3 times in PBS, and incubated with DAPI/PBS for 5 minutes, before a final 2 washes in PBS. Slides were then mounted with 50% glycerol in PBS before imaging. Sections were imaged on a Zeiss LSM 880 Airy Scan Upright laser-scanning confocal microscope with a Plan-Apochromat 20x/0.8 M27 objective. Sections were DAPI stained, labelled with a mouse anti-TH antibody (66334-1-Ig) (ProteinTech, US) and a rabbit antibody against either KCNQ2 (ab22897), KCNQ3 (ab66640) or KCNQ5 (ab66740) (Abcam, UK). For secondary antibodies, 1:200 Donkey anti-mouse Alexa Fluor 555 (A-31570) and 1:200 Donkey anti-rat Alexa Fluor 488 (A-21208) were used (Invitrogen).

### Single cell RNA-sequencing

A pooled single cell suspension of stellate ganglia cells from six animals per strain was prepared via enzymatic dissociation as described under cell culture methods. Following blockade of enzymatic activity via three washes in blocking solution, the cell solution was transferred to phosphate buffered saline. The cell solutions were immediately transferred to ice and transported to the Wellcome Trust Centre for Human Genetics (WTCHG) for scRNAseq via 10x genomics chromium (10x genomics, US) (Single Cell 3’ v3) and Illumina hiseq 4000 (Illumina, US). This approach achieved 66-72K mean reads per cell and a sequencing depth of 53-55% per cell before filtering.

Initial analysis was performed by the WTHCG using the cell ranger pipeline (x10 genomics, US) (Cell ranger, v3.0.2) (Rnor6.0) with default parameters, before the data were exported to Seurat (v3.0) (Stuart *et al*., 2019) and analyzed in house. Cells were excluded in Seurat if the number of counts per cell was less than 4000 or percentage of mitochondrial genes was equal to or less than 0.3. For FindVariableFeatures we used 10000 features and the election method VST. Data was intergrated using 30 dimensions, 30 principle components were using for PCA analysis. UMAP and TSNE, FindNeighbours were ran with 19 dimensions. Findclusters was ran with a resolution of 0.6. Differential expression analysis was performed via MAST (Finak *et al*., 2015) within Seurat. To determine multiple neuron populations for supplementary figure 5, findclusters was ran at a resolution of 10, and neuron-like groups were subset for further analysis. Individual groups were then visually identified based upon clear separation from neighbouring subgroups.

### Electrophysiological Data analysis

Analysis of firing rate data were performed in WinWCP (v5.4.0). M-current deactivation curves were analyzed within Clampfit (v10.7, Molecular Devices, US). Graphs were produced in either Graphpad prism (v8.2.1) or ggplot2 (v3.2.1) and Waffle (v1.0.1) in R (Version 3.5.3). Statistical analysis was performed in GraphPad prism, all relevant datasets were normality tested.

Firing rate was taken as the maximum firing rate elicited by a range of 10 pA current injections between 10-200 pA. Membrane potential was monitored for stability during drug wash in and cells with large jumps in membrane potential were discarded.

Action potential parameters were measured from the first sequential 50 pA current step that induced an action potential. Peak amplitude (mV) was taken as the difference between the average baseline and maximum peak response of the action potential. Action potential upstroke (mV/ms) was taken as the maximum velocity from baseline to the peak amplitude.

Input resistance was calculated based upon a series of hyperpolarizing and depolarizing current injections ranging from −200 to 200 pA in amplitude in 10 pA increments (Spruston and Johnston, 1992). The average value of the final 100 ms was analyzed. As previously classified, small hyperpolarizing pulses were assumed to elicit the least active processes and any points that departed from linearity with these points or contained visible active processes in the final 200 mS of current injection were excluded.

Liquid junction potentials were calculated in JPCalcW (Barry, 1994) in Clampex (v11.0.3) (Molecular Devices, US), where ion availabilities were used instead of concentrations. Free Ca^2+^, ATP, EGTA and Mg^2+^ for internal solutions were estimated via MaxChelator (v8)(Bers, Patton and Nuccitelli, 2010) when relevant. For perforated-patch voltage-clamp and current clamp recordings a Liquid junction potential of 24.3 mV was calculated, without correction for the perforated patch Donnan potential (Horn and Marty, 1988). Whole cell current clamp recordings had an estimated liquid junction potential of −15.7 mV.

### Statistics

All datasets were normality tested, except for firing rate, which was taken as a discontinuous variable and treated as non-parametric data. Statistical analysis and normality tests were performed in graphpad prism (v8.2.1). The specific statistical test applied is stated in the figure legends with statistical significance accepted at p < 0.05 on two tailed tests.

## Acknowledgements

We acknowledge The High-Throughput Genomics Group at the Wellcome Trust Centre for Human Genetics for producing the single cell RNA-sequencing data and for the initial cell-ranger analysis. We acknowledge the British Heart Foundation (RG/17/14/33085) and the Wellcome trust OXION programme and Medical Sciences Doctoral Training Centre, University of Oxford (BST0008Z) for funding this work. NH is supported by a British Heart Foundation Intermediate Fellowship (FS/15/8/3115). We would like to acknowledge Olujimi A. Ajijola M.D. and his team for providing human stellate ganglia for this study.

## Author contributions

H.D performed, interpreted and analyzed all experiments. H.D produced figures. H.D, N.H and D.J.P wrote and edited the manuscript. H.D, N.H and D.J.P designed the project.

## Data availability

The data that support the findings of this study are available from the corresponding authors upon reasonable request. The single cell sequencing dataset generated during this study are available at GEO (GSE144027).

