## Supplementary figure legends for "Novel target identification using single cell sequencing and electrophysiology of cardiac sympathetic neurons in disease"

**Supplementary figure 1** Whole cell recordings of subtype and rheobase confirm that a similar phenotype is present by both electrophysiological configurations. (A) Example traces of Wistar and SHR neurons. (B) Percentage subtype compositions for Wistar and SHR neurons with an increased number of tonic neurons in the SHR (Wistar, n = 38; Phasic 1 n = 8, Phasic 2 n = 19, Tonic n = 11; SHR, n = 52; Phasic 1 n = 13, Phasic 2 n = 13, Tonic n = 26) ($\chi^2 = 1.6043$, df =2, p-value = 0.45).

**Supplementary figure 2** Top changes in neuronal cluster gene expression between Wistar and SHR strains (relative to Wistar) as determined by Single cell RNA-sequencing. (A) Top 50 genes that are significantly increased in the SHR. (B) Top 50 genes that are significantly decreased in the SHR. (C) Top 50 genes expressed independent of strain in Wistar and SHR neurons. (D) Channel encoding genes assessed for comparison of genes involved in firing rate.

**Supplementary figure 3** Cluster expression of channels with low expression levels or without specific inhibitors. Expression of ion channel genes listed in Supplementary figure 2D, but not pharmacologically tested within the study are shown graphically here for future study of stellate ganglia cell populations. Cacna1i, Kcnb2 and Kcnd2 were not detected in any cell type and are therefore not represented in either figure.

**Supplementary figure 4** Classic markers for identified clusters in the stellate ganglia. (A) Smooth muscle actin (Acta2), Myosin heavy chain 11 (Myh11) and Transgelin (Tagln) were used as markers for the Vascular smooth muscle cluster. (B) CD45 (Ptprc) was used as a pan-immune cell marker. (C) Glial-fibrillary acidic protein (Gfap), Myelin protein zero (Mpz) and S100 calcium binding protein B (S100b) were used as glial markers. (D) Cadherin 5 (Cdh5), Intercellular Adhesion Molecule 2 (Icam2), Kinase Insert Domain Receptor (Kdr), Selectin E (Sele), and Selectin P (Selp) were used as Endothelial cell markers. (E) Decorin (Dcn), Fibroblast Activation Protein Alpha (Fap), Lumican (Lum), Matrix Gla Protein (Mgp) and SRY-Box Transcription Factor 9 (Sox9) were used as Fibroblast markers.

**Supplementary figure 5** Unbiased markers for major cell clusters found in the stellate ganglia as detected by Seurat analysis. (A) Top 30 markers for the immune cell cluster. (B) Top 30 markers for the Vascular smooth muscle cell cluster. (C) Top 30 markers for the glial cell cluster. (D) Top 30 markers for the Fibroblast cell cluster. (E) Top 30 markers for the endothelial cell cluster. (F) Top 30 markers for sympathetic neuron clusters.

**Supplementary figure 6** Evidence of multiple populations of sympathetic neurons. (A) Visually identified cell clusters of potential sympathetic neurons. (B) Expression of key transcripts for noradrenaline synthesis and breakdown are shown for each neuron cluster. Of these groups only Neurons 1 and Neurons 2 appear to adequately express the full noradrenaline synthesis pathway. (C) Sympathetic Neurons 1 are shown to be have low expression of CHRM2 and NPY, two physiologically important genes, in contrast to high expression in Sympathetic Neurons 2. For reference, these groups are termed Type A sympathetic neurons (Sympathetic Neurons 1), and type B sympathetic neurons (Sympathetic Neurons 2). Importantly, we also demonstrate that both groups have similar ion channel profiles, with no significant differences observed between groups.

**Supplementary figure 7** The effect of M-current on other parameters, the expression of M-current per SHR firing rate subtype and Immunohistochemistry for M-current. (A) M-current inhibition by 3 µM XE-991 caused a depolarization of Wistar neuron resting membrane potential (Mean ± SEM) (Control, -72.59 ± 0.97 mV; 3 µM XE-991, -67.07 ± 1.08 mV) (Paired t-test, n = 17, $p < 0.0001$). (B) M-current inhibition by 30 µM Linopirdine also causes a depolarization of resting membrane potential (Mean ± SEM) (Control, -73.01 ± 1.37 mV; 30 µM Linopirdine, -69.29 ± 1.72 mV) (Paired t-test, n = 11, p = 0.0003). (C) M-current activation by retigabine causes a hyperpolarization of resting membrane potential at higher doses (Mean ± SEM) (Control, -70.27 ± 1.39 mV; 3 µM, -83.63 ± 1.67 mV; 10 µM, -87.69 ± 1.53 mV; 30 µM, -88.45 ± 1.91 mV) (One-way ANOVA, n = 12, $p < 0.0001$) (Dunnett's multiple comparisons; Control vs 3 µM, $p < 0.0001$; Control vs 10 µM, $p < 0.0001$; Control vs 30 µM, $p < 0.0001$). (D) M-current inhibition by 3 µM XE-991 decreased Wistar neuron Rheobase (Median) (Control, 50 pA; 3 µM XE-991, 20 pA) (Wilcoxon test, n = 17, $p < 0.0001$).(E) M-current inhibition by 30 µM Linopirdine decreased Wistar Rheobase (Median) (Control, 55 pA; 30 µM Linopirdine, 35 pA)

(Wilcoxon test, n = 10, p = 0.0039). (F) M-current activation by 3 μM retigabine increased SHR rheobase (Median) (Control, 30 pA; 3 μM retigabine, 110 pA) (Wilcoxon test, n = 9, p = 0.0039). (G) Immunohistochemistry showing M-current subunit protein in TH-positive neurons in cryosections of 5-6 week old Wistar stellate ganglia (KCNQ2, KCNQ3 and KCNQ5).

**Supplementary figure 8** Additional data in support of figure 6. (A) Low dose TTX (10 nM) significantly decreased SHR action potential upstroke velocity (Mean ± SEM) (Control 143.2 ± 12.73 mV/ms, 10 nM 91.09 ± 11.14 mV/ms) (Paired T-test, n = 13, p = 0.0003). (B) Low dose TTX (10 nM) significantly decreased SHR action potential amplitude (Median) (Control 102.5 mV, 10 nM 85.8 mV) (Wilcoxon test, n = 13, p = 0.0046). (C) No significant difference was observed between Wistar and SHR action potential upstroke velocity (Median ± IQR) (Wistar 64.72 mV/ms, n = 38; SHR 70.54 mV/ms, n = 52) (Mann-Whitney test, p = 0.067). (D) No significant difference was observed between Wistar and SHR action potential amplitude (Median ± IQR) (Wistar, 75.74 mV, n = 37; SHR, 79.39 mV, n = 52) (Mann-Whitney test; p = 0.27). (E) 100 nM 4,9-Anhydrotetrotoxin decreased action potential upstroke velocity (Mean ± SEM) (Control 85.94 ± 16.07 mV/ms, 4,9-Anhydrotetrotoxin 61.54 ± 12.99 mV/ms) (Paired t-test, n = 12 , p = 0.023). (F) 300 nM TTX prevented firing in all tested SHR neurons (Median) (Control 2 Hz, 300 nM TTX 0 Hz) (Wilcoxon test, n = 9, p = 0.0039). (G) Dependence of firing rate on membrane potential was determined in SHR neurons by applying a pre-pulse in the range -10 to -100 pA for 1 second before applying a 150 pA positive current injection to elicit cell firing. No difference was found (Friedman test, n = 26, p = 0.165).

**Supplementary figure 9** The previously described inhibitory effect of $Na_v$1.8 inhibition on stellate ganglia function in vivo likely occurs through non-specific sodium channel inhibition. (A) $Na_v$1.8 inhibitors 100 nM A803467 and 300 nM A887826 significantly decrease maximal firing rate of SHR stellate ganglia neurons (Median) (Control, 6 Hz; 100 nM A803467, 1.5 Hz; n = 10, p = 0.002; Control, 8 Hz; 300 nM A887826, 2 Hz) (Wilcoxon test, n = 11, p = 0.002). (B) $Na_v$1.8 inhibitor 1 μM PF04885614 does not affect maximal firing rate of SHR stellate ganglia neurons (Median) (Control, 16 Hz;

PF04885614, 19 Hz) (Wilcoxon test, n = 9, p = 0.17). (C) $Na_v1.8$ inhibitor 100 nM A803467 did not affect resting membrane potential (Mean ± SEM) (Control, -63.99 ± 2.9 mV; A803467, -64.34 ± 4.49 mV) (Paired t-test, n = 10, p = 0.88). (D) $Na_v1.8$ inhibitor 300 nM A887826 did not affect resting membrane potential (Mean ± SEM) (Control, -62.61 ± 2.75 mV; A887826, 65.43 ± 2.48 mV) (Paired t-test, n = 11, p = 0.21). (E) $Na_v1.8$ inhibitor 1 µM PF04885614 did not affect resting membrane potential (Mean ± SEM) (Control, -61.97 ± 2.12 mV; PF04885614, -61.63 ± 3.93 mV) (Paired t-test, n = 9, p = 0.92).

**Supplementary figure 10** Analysis of the relationship between cell capacitance, days in vitro (DIV) and maximum firing rate in either strain as measured by perforated patch clamp. (A) The number of days in does not affect the maximum neuronal firing rate in Wistar neurons (Median) (DIV1, 2 Hz, n = 19; DIV2, 2 Hz, n = 14; DIV3, 1.5 Hz, n = 28; DIV4, 1 Hz, n = 5) (Kruskal-Wallis test, n = 66, p = 0.48). (B) There is no clear relationship between cell capacitance and firing rate in Wistar neurons. (C) The number of days in does not affect the maximum neuronal firing rate in SHR neurons (Median) (DIV1, 5.5 Hz, n = 18; DIV2, 9 Hz, n = 22; DIV3, 9.5 Hz, n = 18; DIV4, 4 Hz, n = 10; DIV5, 8.5 Hz, n = 2) (Kruskal-Wallis test, n = 70, p = 0.35). (D) There is no clear relationship between cell capacitance and firing rate in SHR neurons. (E) There is not a significant difference in the number of days in vitro between strains (Median) (Wistar, 2.5 days, n = 66; SHR, 2 days, n = 70) (Mann-Whitney test, p = 0.83). (F) There is no significant difference in capacitance between strains, although there is a non-significant trend (Median) (Wistar, 26.74 pF, n = 66; SHR, 23.35 pF, n = 70) (Mann-Whitney, p = 0.27).
