## Supplementary figures for "Novel target identification using single cell sequencing and electrophysiology of cardiac sympathetic neurons in disease"

S1

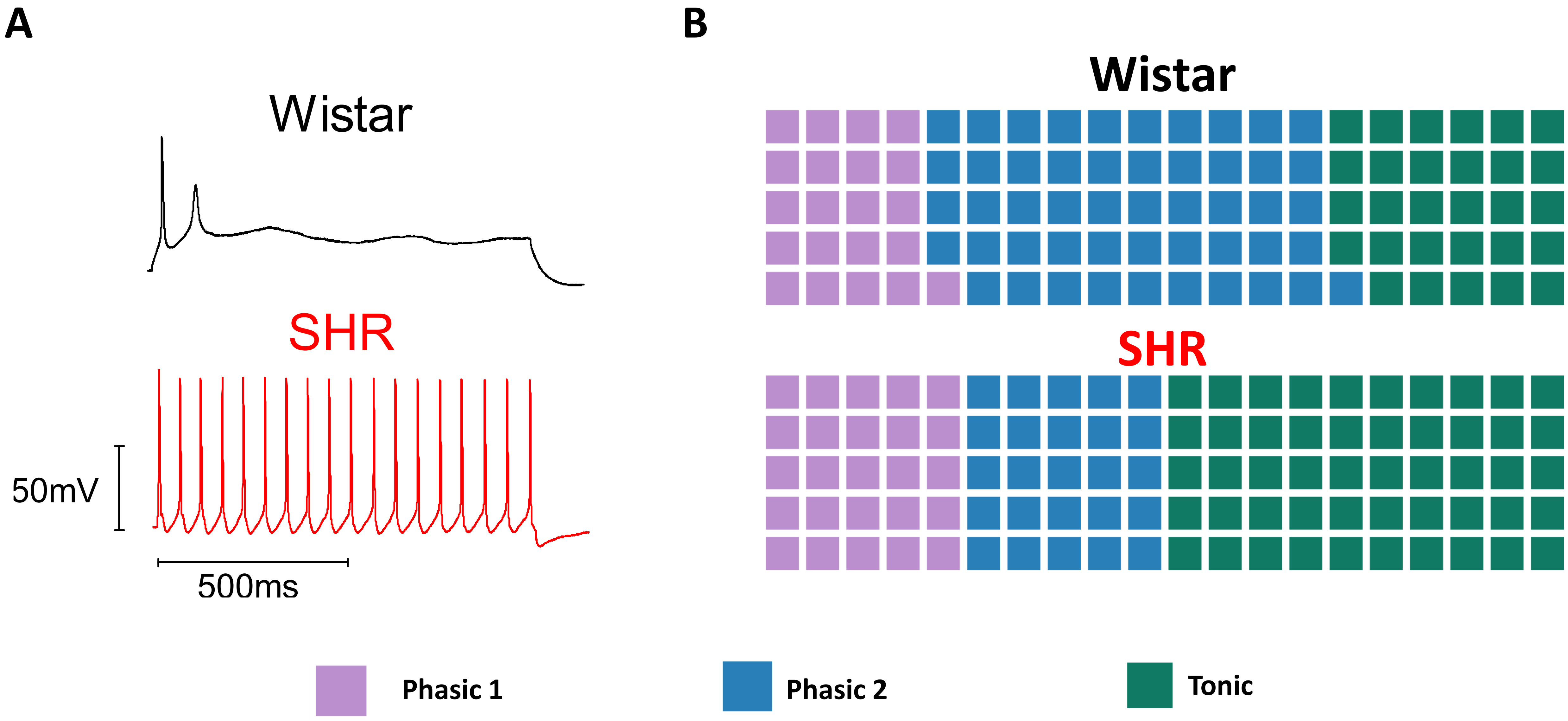

# S2

### Neurons

**A**

#### Increased expression

| Gene | Log2Fold change | Adjusted P value |
|---|---|---|
| Jund | 1.5082516 | 4.367859e-101 |
| S100a6 | 1.0578275 | 2.450488e-09 |
| LOC100364435 | 0.7454090 | 5.585682e-14 |
| Lgals3 | 0.7439313 | 1.889339e-21 |
| Rgs10 | 0.6371928 | 2.374208e-20 |
| Cox7c | 0.6360328 | 1.545634e-14 |
| Ifi27l2b | 0.5969734 | 1.413566e-10 |
| Rps25 | 0.5790300 | 2.864862e-22 |
| Nbl1 | 0.5747978 | 6.333336e-11 |
| Lgals1 | 0.5714637 | 1.392294e-13 |
| RT1-A1 | 0.5460656 | 7.680405e-20 |
| Atp5f1e | 0.5318127 | 1.876125e-08 |
| Tmsb4x | 0.5265573 | 3.358830e-05 |
| Lix1 | 0.5142321 | 3.978586e-15 |
| Cops9 | 0.4927932 | 1.323195e-15 |
| Gsta4 | 0.4855133 | 3.403365e-50 |
| Gabarap | 0.4794821 | 8.050229e-08 |
| Sncg | 0.4733403 | 9.515786e-06 |
| Lynx1 | 0.4652957 | 2.303662e-13 |
| Fxyd6 | 0.4546705 | 1.184943e-11 |
| Gap43 | 0.4534140 | 1.044454e-11 |
| LOC100359583 | 0.4523299 | 6.131520e-09 |
| Ube2s | 0.4504232 | 7.396337e-23 |
| LOC103689961 | 0.4501770 | 5.011713e-04 |
| Rps15a | 0.4469772 | 3.038277e-04 |
| Nrsn1 | 0.4431640 | 3.459381e-19 |
| Cd24 | 0.4429852 | 4.084656e-10 |
| Pfn1 | 0.4382201 | 5.952396e-06 |
| Clic3 | 0.4381656 | 2.269674e-21 |
| Id1 | 0.4362874 | 3.333847e-09 |
| S100a11 | 0.4360989 | 3.040743e-07 |
| Cstb | 0.4340093 | 1.223860e-12 |
| Dynll1 | 0.4320592 | 6.394903e-09 |
| Rpl6 | 0.4313981 | 2.959553e-07 |
| Mt3 | 0.4196665 | 1.086267e-03 |
| Ywhah | 0.4167402 | 9.640197e-09 |
| S100a10 | 0.4156335 | 5.061056e-08 |
| Crip1 | 0.4131412 | 1.000000e+00 |
| Cox6c | 0.4121257 | 2.138002e-05 |
| RGD1564664 | 0.4105464 | 6.250173e-09 |
| Cox8a | 0.4105403 | 2.700923e-03 |
| Ctxn1 | 0.4092833 | 2.431107e-07 |
| Tppp3 | 0.4087822 | 1.062926e-12 |
| Fabp5 | 0.4030572 | 4.743981e-08 |
| Atp5mc2 | 0.4022625 | 1.413702e-05 |
| Rps27a.1 | 0.4015203 | 1.378063e-02 |
| Uqcrq | 0.3967584 | 1.291659e-07 |
| Ndufb2 | 0.3958326 | 9.469155e-06 |
| LOC687780 | 0.3952969 | 4.839876e-04 |
| Sncb | 0.3943067 | 5.081632e-09 |
| Rps4x.1 | 0.3931921 | 1.000000e+00 |
| Ost4 | 0.3926721 | 8.414767e-07 |
| Fkbp1b | 0.3924926 | 1.496808e-04 |
| Rpl41 | 0.3922107 | 9.844968e-04 |
| Rpl24 | 0.3918698 | 5.646911e-02 |
| Elob | 0.3912504 | 9.582211e-06 |
| Rpl10 | 0.3908021 | 9.266765e-06 |
| Id3 | 0.3898860 | 2.287913e-09 |
| Timm8b | 0.3896933 | 1.247257e-06 |
| Atp5pf | 0.3882082 | 1.176928e-06 |
| Rpsa | 0.3863797 | 6.523119e-06 |
| Ptms | 0.3860194 | 5.308388e-10 |
| Rps24 | 0.3823999 | 1.109389e-04 |

**B**

#### Decreased expression

| Gene | Log2Fold change | Adjusted P value |
|---|---|---|
| AC134224.1 | -1.1919235 | 7.390003e-14 |
| AC134224.3 | -1.1243569 | 2.337894e-01 |
| Rsrp1 | -1.0110946 | 8.436000e-07 |
| Cd9 | -1.0105276 | 2.551646e-14 |
| Pcp4 | -0.8881339 | 2.445960e-21 |
| Avil | -0.8147849 | 1.833540e-32 |
| Apoe | -0.8061488 | 1.826096e-01 |
| Snhg11 | -0.7870100 | 1.000000e+00 |
| Clasrp | -0.7413731 | 4.021941e-03 |
| Gria2 | -0.7367756 | 2.002588e-10 |
| Snrnp70 | -0.7127888 | 4.602979e-02 |
| AABR07043389.1 | -0.7011961 | 7.905533e-25 |
| Rock1 | -0.6985003 | 1.892195e-04 |
| Tnpo1 | -0.6799992 | 1.659938e-15 |
| Mcf2l | -0.6690244 | 1.769559e-07 |
| Slc12a3 | -0.6554360 | 2.632086e-24 |
| Xkr6 | -0.6535802 | 3.527137e-01 |
| Insrr | -0.6426622 | 6.661967e-04 |
| AC134224.2 | -0.6362384 | 1.233835e-04 |
| Kifc2 | -0.6291232 | 3.462901e-03 |
| Alcam | -0.6282179 | 2.411339e-07 |
| Macf1 | -0.6187952 | 1.524047e-14 |
| Ddc | -0.5950884 | 2.365693e-18 |
| Rbfox1 | -0.5945057 | 5.243143e-03 |
| Pnisr | -0.5908631 | 6.034754e-03 |
| Leng8 | -0.5850844 | 1.022591e-03 |
| Aqp1 | -0.5821055 | 4.037932e-08 |
| Ssbp4 | -0.5752207 | 2.196309e-01 |
| Abca7 | -0.5728495 | 2.284495e-03 |
| Epha5 | -0.5728401 | 8.027044e-07 |
| Ogt | -0.5657906 | 7.585591e-07 |
| Srrm2 | -0.5603236 | 1.537572e-02 |
| Srsf2 | -0.5566520 | 3.052943e-04 |
| Zbtb20 | -0.5551126 | 3.212383e-05 |
| Stk38 | -0.5509250 | 1.375099e-06 |
| Carmil3 | -0.5479098 | 4.352376e-03 |
| Lss | -0.5475808 | 1.238291e-03 |
| Agrn | -0.5446013 | 3.798112e-04 |
| Prkg2 | -0.5419457 | 9.840020e-18 |
| Ddx39b | -0.5372624 | 6.367377e-05 |
| Brinp2 | -0.5350721 | 4.241865e-05 |
| Pnn | -0.5290578 | 6.977921e-02 |
| Pcmtd2 | -0.5265765 | 3.677216e-03 |
| Sparc | -0.5206209 | 1.000000e+00 |
| Pabpn1 | -0.5186682 | 9.728370e-01 |
| Vav2 | -0.5151115 | 2.180735e-06 |
| Pclo | -0.5125366 | 3.626956e-08 |
| Arhgap21 | -0.5025987 | 7.737147e-11 |
| P3h3 | -0.5017257 | 5.764083e-10 |
| Trafd1 | -0.4986911 | 4.079081e-03 |
| Zcchc7 | -0.4942523 | 1.663439e-01 |
| Nfat5 | -0.4886901 | 4.319940e-05 |
| Atp13a2 | -0.4856006 | 2.457154e-08 |
| Taf1d | -0.4822939 | 1.000000e+00 |
| Luc7l3 | -0.4804302 | 3.571888e-03 |
| Atxn2l | -0.4792462 | 6.461217e-02 |
| Cacna1b | -0.4773798 | 1.671714e-04 |
| Chl1 | -0.4741254 | 9.661983e-10 |
| Abca8a | -0.4734990 | 1.000000e+00 |
| RT1-CE4 | -0.4724680 | 3.738463e-21 |
| Ank2 | -0.4707748 | 2.647609e-12 |
| Dkc1 | -0.4665386 | 1.870979e-02 |
| Clk1 | -0.4629965 | 3.111367e-02 |

**C**

#### Top expressed

| Gene | Wistar Reads |
|---|---|
| Mt-cyb | 5.441520 |
| Mt-co1 | 5.067902 |
| Mt-nd2 | 4.896349 |
| Mt-nd1 | 4.574465 |
| Mt-nd4 | 4.544456 |
| Tuba1a | 4.350063 |
| LOC100364435 | 4.156522 |
| Prph | 4.076758 |
| AC134224.1 | 3.993742 |
| Stmn2 | 3.834467 |
| Tubb3 | 3.824851 |
| Map1b | 3.812369 |
| Sncg | 3.809325 |
| Tmsb4x | 3.784265 |
| Hsp90ab1 | 3.410114 |
| Snhg11 | 3.294435 |
| Atp6v0c | 3.286958 |
| Uchl1 | 3.257605 |
| Atp1a1 | 3.224183 |
| Actg1 | 3.214234 |
| Syt1 | 3.194060 |
| Tubb5 | 3.174731 |
| Rtn1 | 3.150431 |
| Stmn3 | 3.148980 |
| Tubb2a | 3.106304 |
| Fth1 | 3.086398 |
| Atp1b1 | 3.023032 |
| Tubb2b | 3.013394 |
| Nefl | 3.004773 |
| Snap25 | 2.914318 |
| S100a6 | 2.890331 |
| Zwint | 2.872957 |
| Syt4 | 2.869539 |
| Stmn1 | 2.865590 |
| Ndrg4 | 2.860553 |
| Cd9 | 2.846434 |
| Elavl2 | 2.840360 |
| Ndfip1 | 2.839060 |
| Slc6a2 | 2.825227 |
| Basp1 | 2.824477 |
| Rtn3 | 2.799926 |
| Calm1 | 2.788565 |
| Ywhah | 2.779392 |
| Reep5 | 2.768612 |
| Cyb561 | 2.765191 |
| Ntrk1 | 2.736131 |
| Dst | 2.726988 |
| Tspan8 | 2.718403 |
| Cfl1 | 2.716252 |
| App | 2.709063 |
| Cd24 | 2.707676 |
| Nsg1 | 2.702262 |
| LOC103692716 | 2.695446 |
| Rtn4 | 2.687459 |
| Gap43 | 2.659190 |
| Bcat1 | 2.658582 |
| LOC108348172.1 | 2.656803 |
| Aldoa | 2.631778 |
| Lgals1 | 2.622184 |

**D**

#### Channels assessed

| Gene | Channel |
|---|---|
| Scn1a | $Na_v1.1$ |
| Scn2a | $Na_v1.2$ |
| Scn3a | $Na_v1.3$ |
| Scn4a | $Na_v1.4$ |
| Scn5a | $Na_v1.5$ |
| Scn7a | $Na_v2.1$ |
| Scn8a | $Na_v1.6$ |
| Scn9a | $Na_v1.7$ |
| Scn10a | $Na_v1.8$ |
| Scn11a | $Na_v1.9$ |
| Hcn1 | HCN1 |
| Hcn2 | HCN2 |
| Hcn3 | HCN3 |
| Hcn4 | HCN4 |
| Ano1 | TMEM16A |
| Ano2 | TMEM16B |
| Cacna1c | $Ca_v1.2$ |
| Cacna1d | $Ca_v1.3$ |
| Cacna1a | $Ca_v2.1$ |
| Cacna1b | $Ca_v2.2$ |
| Cacna1g | $Ca_v3.1$ |
| Cacna1h | $Ca_v3.2$ |
| Cacna1i | $Ca_v3.3$ |
| Kcnt1 | SLACK |
| Kcnt2 | SLICK |
| Kcnma1 | BK($K_{Ca}1.1$) |
| Kcnn1 | SK1($K_{Ca}2.1$) |
| Kcnn2 | SK2($K_{Ca}2.2$) |
| Kcnn3 | SK2($K_{Ca}2.3$) |
| Kcnn4 | IK($K_{Ca}3.1$) |
| Kcna1 | $K_v1.1$ |
| Kcna2 | $K_v1.2$ |
| Kcna3 | $K_v1.3$ |
| Kcna4 | A-type ($K_v1.4$) |
| Kcna5 | $K_v1.5$ |
| Kcna6 | $K_v1.6$ |
| Kcna7 | $K_v1.7$ |
| Kcnb1 | $K_v2.1$ |
| Kcnb2 | $K_v2.2$ |
| Kcnc1 | $K_v3.1$ |
| Kcnc2 | $K_v3.2$ |
| Kcnc3 | A-type ($K_v3.3$) |
| Kcnc4 | A-type ($K_v3.4$) |
| Kcnd1 | A-type ($K_v4.1$) |
| Kcnd2 | A-type($K_v4.2$) |
| Kcnd3 | A-type($K_v4.3$) |
| Kcnf1 | $K_v5.1$ |
| Kcng1 | $K_v6.1$ |
| Kcng2 | $K_v6.2$ |
| Kcng4 | $K_v6.3$ |
| Kcng3 | $K_v6.4$ |
| Kcnq2 | $K_v7.2$ |
| Kcnq3 | $K_v7.3$ |
| Kcnq4 | $K_v7.4$ |
| Kcnq5 | $K_v7.5$ |
| Kcnk1 | $K_{2P}1.1$ |
| Kcnk2 | $K_{2P}2.1$ |
| Kcnk3 | $K_{2P}3.1$ |
| Kcnk4 | $K_{2P}4.1$ |
| Kcnk5 | $K_{2P}5.1$ |
| Kcnk6 | $K_{2P}6.1$ |
| Kcnk7 | $K_{2P}7.1$ |

S3

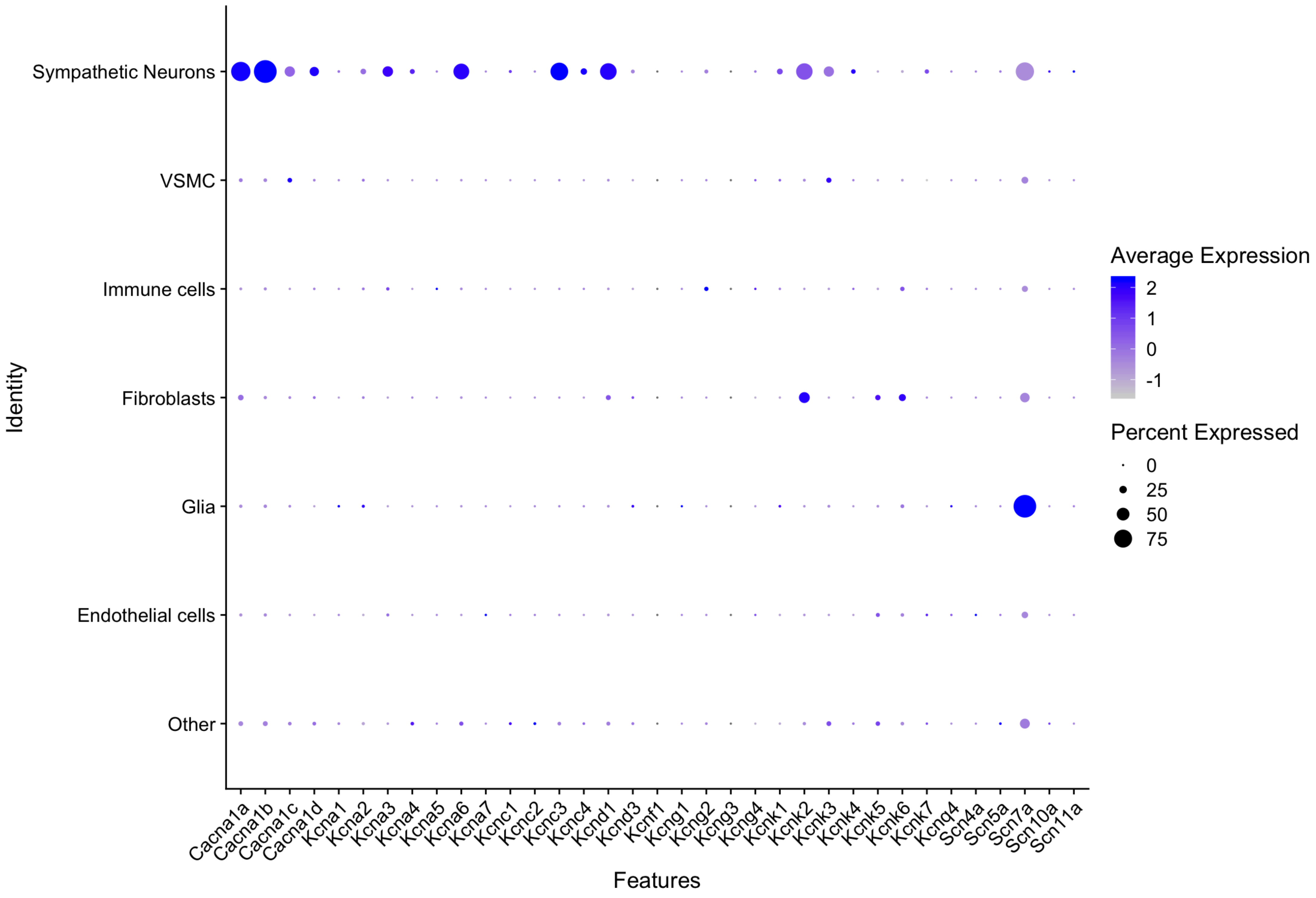

Sympathetic Neurons
VSMC
Immune cells
Fibroblasts
Glia
Endothelial cells
Other
Identity
Cacna1a
Cacna1b
Cacna1c
Cacna1d
Kcna1
Kcna2
Kcna3
Kcna4
Kcna5
Kcna6
Kcna7
Kcnc1
Kcnc2
Kcnc3
Kcnc4
Kcnd1
Kcnd3
Kcnf1
Kcng1
Kcng2
Kcng3
Kcng4
Kcnk1
Kcnk2
Kcnk3
Kcnk4
Kcnk5
Kcnk6
Kcnk7
Kcnq4
Scn4a
Scn5a
Scn7a
Scn10a
Scn11a
Features
Average Expression
2
1
0
-1
Percent Expressed
0
25
50
75

S4

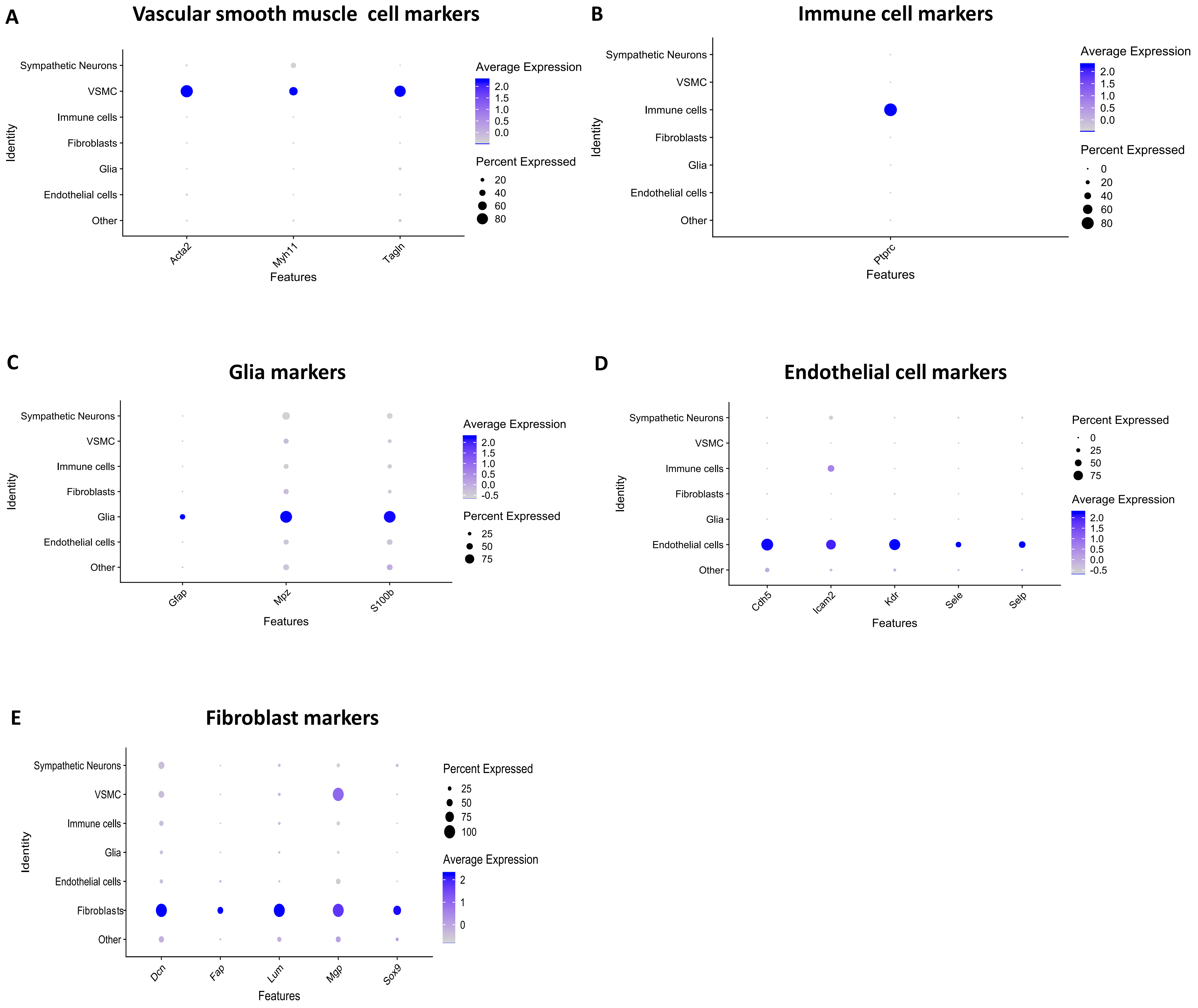
A
Vascular smooth muscle cell markers
Sympathetic Neurons
VSMC
Immune cells
Fibroblasts
Glia
Endothelial cells
Other
Identity
Acta2
Myh11
Tagln
Features
Average Expression
2.0
1.5
1.0
0.5
0.0
Percent Expressed
20
40
60
80
B
Immune cell markers
Sympathetic Neurons
VSMC
Immune cells
Fibroblasts
Glia
Endothelial cells
Other
Identity
Ptprc
Features
Average Expression
2.0
1.5
1.0
0.5
0.0
Percent Expressed
0
20
40
60
80
C
Glia markers
Sympathetic Neurons
VSMC
Immune cells
Fibroblasts
Glia
Endothelial cells
Other
Identity
Gfap
Mpz
S100b
Features
Average Expression
2.0
1.5
1.0
0.5
0.0
-0.5
Percent Expressed
25
50
75
D
Endothelial cell markers
Sympathetic Neurons
VSMC
Immune cells
Fibroblasts
Glia
Endothelial cells
Other
Identity
Cdh5
Icam2
Kdr
Sele
Selp
Features
Percent Expressed
0
25
50
75
Average Expression
2.0
1.5
1.0
0.5
0.0
-0.5
E
Fibroblast markers
Sympathetic Neurons
VSMC
Immune cells
Glia
Endothelial cells
Fibroblasts
Other
Identity
Dcn
Fap
Lum
Mgp
Sox9
Features
Percent Expressed
25
50
75
100
Average Expression
2
1
0

# S5

## A

#### Immune cell Markers

| Gene | Log2Fold change | Percentage 1 | Percentage 2 | Adjusted P value |
|---|---|---|---|---|
| C1qa | 3.5180842 | 0.877 | 0.046 | 0.000000e+00 |
| Cxcl2 | 3.6511131 | 0.743 | 0.027 | 0.000000e+00 |
| C1qb | 3.4324098 | 0.823 | 0.038 | 0.000000e+00 |
| C1qc | 3.3461027 | 0.827 | 0.037 | 0.000000e+00 |
| Tyrobp | 2.9095431 | 0.937 | 0.042 | 0.000000e+00 |
| Fcer1g | 2.8219276 | 0.930 | 0.037 | 0.000000e+00 |
| Cfh | 2.8945865 | 0.897 | 0.032 | 0.000000e+00 |
| Aif1 | 2.6733969 | 0.910 | 0.065 | 0.000000e+00 |
| Il1b | 3.0137275 | 0.760 | 0.023 | 0.000000e+00 |
| Bcl2a1 | 2.6581615 | 0.850 | 0.023 | 0.000000e+00 |
| Cd83 | 2.7810403 | 0.883 | 0.018 | 0.000000e+00 |
| Mrc1 | 2.4344799 | 0.867 | 0.021 | 0.000000e+00 |
| Ccl6 | 2.3750514 | 0.840 | 0.015 | 0.000000e+00 |
| Pf4 | 2.1622164 | 0.780 | 0.014 | 0.000000e+00 |
| Laptm5 | 2.0547600 | 0.993 | 0.058 | 0.000000e+00 |
| Cybb | 2.3030469 | 0.893 | 0.014 | 0.000000e+00 |
| Csf1r | 2.2658411 | 0.880 | 0.016 | 0.000000e+00 |
| Clec10a | 2.2311328 | 0.870 | 0.018 | 0.000000e+00 |
| Rgs1 | 1.8959000 | 0.790 | 0.012 | 0.000000e+00 |
| Cfd | 1.4389330 | 0.637 | 0.009 | 0.000000e+00 |

## B

#### Vascular smooth muscle cell Markers

| Gene | Log2Fold change | Percentage 1 | Percentage 2 | Adjusted P value |
|---|---|---|---|---|
| Acta2 | 3.4346256 | 0.845 | 0.022 | 0.000000e+00 |
| Tagln | 3.2154856 | 0.764 | 0.078 | 9.542926e-145 |
| RGD1564664 | 2.9895026 | 0.991 | 0.253 | 1.159849e-102 |
| Myh11 | 2.8686134 | 0.627 | 0.054 | 4.842476e-133 |
| Rasl11a | 2.6877606 | 0.891 | 0.055 | 8.564578e-253 |
| Mustn1 | 2.5042547 | 0.855 | 0.117 | 1.658917e-132 |
| Tpm2 | 2.4530870 | 0.882 | 0.275 | 2.984776e-73 |
| Myl9 | 2.4404292 | 0.891 | 0.239 | 4.692441e-85 |
| Mgp | 2.3238439 | 1.000 | 0.211 | 1.751286e-112 |
| Des | 2.1621336 | 0.818 | 0.022 | 0.000000e+00 |
| Vtn | 2.0786987 | 0.855 | 0.060 | 3.201642e-213 |
| Cox4i2 | 1.9366285 | 0.918 | 0.196 | 9.119496e-104 |
| Rgs5 | 1.8996050 | 0.736 | 0.022 | 0.000000e+00 |
| Tpm1 | 1.8941308 | 0.918 | 0.575 | 5.534833e-44 |
| Fabp4 | 1.8823574 | 0.718 | 0.025 | 2.387438e-280 |
| Adamts1 | 1.8798563 | 0.700 | 0.179 | 1.850121e-52 |
| Mylk | 1.8776962 | 0.836 | 0.091 | 4.514715e-153 |
| Npy1r | 1.8622683 | 0.855 | 0.031 | 0.000000e+00 |
| Ndufa4l2 | 1.8418009 | 0.918 | 0.097 | 8.715348e-177 |
| Hopx | 1.7450186 | 0.864 | 0.050 | 5.282456e-251 |

## C

#### Glia cell Markers

| Gene | Log2Fold change | Percentage 1 | Percentage 2 | Adjusted P value |
|---|---|---|---|---|
| Dbi | 2.0501341 | 1.000 | 0.976 | 0.000000e+00 |
| Fxyd1 | 2.0230451 | 0.998 | 0.679 | 0.000000e+00 |
| Sostdc1 | 1.9505253 | 0.968 | 0.292 | 0.000000e+00 |
| Scn7a | 1.8488933 | 0.990 | 0.387 | 0.000000e+00 |
| Cdh19 | 1.7355892 | 0.980 | 0.265 | 0.000000e+00 |
| Vwa1 | 1.6739017 | 0.992 | 0.472 | 0.000000e+00 |
| Abca8a | 1.6606951 | 0.994 | 0.604 | 0.000000e+00 |
| Sfrp5 | 1.6566244 | 0.952 | 0.317 | 0.000000e+00 |
| Art3 | 1.5968683 | 0.992 | 0.335 | 0.000000e+00 |
| S100b | 1.5535275 | 0.982 | 0.475 | 0.000000e+00 |
| LOC108348061 | 1.5519136 | 0.860 | 0.415 | 0.000000e+00 |
| Gpr37l1 | 1.5491717 | 0.945 | 0.199 | 0.000000e+00 |
| Cnn3 | 1.5430965 | 0.995 | 0.760 | 0.000000e+00 |
| Lgi4 | 1.5341173 | 0.979 | 0.311 | 0.000000e+00 |
| Col28a1 | 1.5185676 | 0.912 | 0.200 | 0.000000e+00 |
| Pdlim4 | 1.5179088 | 0.966 | 0.323 | 0.000000e+00 |
| Tmod2 | 1.5000766 | 0.929 | 0.392 | 0.000000e+00 |
| Gpm6b | 1.4857464 | 0.972 | 0.330 | 0.000000e+00 |
| Rarres2 | 1.4785534 | 0.986 | 0.456 | 0.000000e+00 |
| Egfl8 | 1.4631824 | 0.970 | 0.365 | 0.000000e+00 |

## D

#### Fibroblast Markers

| Gene | Log2Fold change | Percentage 1 | Percentage 2 | Adjusted P Value |
|---|---|---|---|---|
| Apod | 5.0702161 | 0.972 | 0.444 | 8.531948e-70 |
| Dcn | 4.4300866 | 1.000 | 0.257 | 1.504777e-106 |
| Myoc | 4.2646390 | 0.991 | 0.091 | 5.961128e-216 |
| Lum | 3.8051183 | 1.000 | 0.113 | 1.256168e-188 |
| Gsn | 3.0628327 | 1.000 | 0.824 | 3.071820e-63 |
| Thbs4 | 2.9821255 | 1.000 | 0.044 | 0.000000e+00 |
| Col3a1 | 2.9581712 | 1.000 | 0.846 | 9.210442e-65 |
| Mgp | 2.8940880 | 0.991 | 0.212 | 2.892069e-110 |
| Igfbp5 | 2.6937379 | 0.755 | 0.328 | 2.660498e-33 |
| Col1a1 | 2.6731355 | 1.000 | 0.823 | 4.557282e-63 |
| Col15a1 | 2.5881164 | 1.000 | 0.545 | 6.192017e-70 |
| Crispld2 | 2.5701496 | 0.943 | 0.042 | 0.000000e+00 |
| Fn1 | 2.5640687 | 1.000 | 0.250 | 3.140104e-102 |
| Gpc3 | 2.5392989 | 0.972 | 0.024 | 0.000000e+00 |
| Smoc2 | 2.4793013 | 0.934 | 0.027 | 0.000000e+00 |
| Serpinf1 | 2.4357652 | 0.972 | 0.066 | 6.852849e-261 |
| Pcolce | 2.4270370 | 1.000 | 0.151 | 6.559569e-153 |
| Mmp2 | 2.3279613 | 1.000 | 0.120 | 2.809839e-179 |
| Aebp1 | 2.2966367 | 0.981 | 0.103 | 3.657190e-192 |
| Igfbp6 | 2.2607160 | 0.830 | 0.096 | 1.724846e-134 |

## E

#### Endothelial cell Markers

| Gene | Log2Fold change | Percentage 1 | Percentage 2 | Adjusted P Value |
|---|---|---|---|---|
| Plvap | 2.6699906 | 0.976 | 0.030 | 0.000000e+00 |
| Aqp1 | 2.5668906 | 0.961 | 0.146 | 0.000000e+00 |
| Rgs16 | 2.5153660 | 0.821 | 0.101 | 0.000000e+00 |
| Sele | 2.4944279 | 0.374 | 0.014 | 7.421128e-235 |
| Flt1 | 2.3509513 | 0.966 | 0.019 | 0.000000e+00 |
| Slco1a4 | 2.1993135 | 0.918 | 0.029 | 0.000000e+00 |
| Selp | 2.1762584 | 0.466 | 0.018 | 4.578829e-295 |
| Emcn | 2.1625471 | 0.978 | 0.038 | 0.000000e+00 |
| Abcg2 | 2.1382139 | 0.913 | 0.041 | 0.000000e+00 |
| Cldn5 | 2.1186817 | 0.787 | 0.024 | 0.000000e+00 |
| Cyyr1 | 2.1040015 | 0.971 | 0.028 | 0.000000e+00 |
| Fam110d | 2.0985356 | 0.886 | 0.022 | 0.000000e+00 |
| Rnd1 | 2.0177000 | 0.659 | 0.071 | 2.776598e-287 |
| Cav1 | 2.0059116 | 0.957 | 0.087 | 0.000000e+00 |
| Cxcl12 | 1.9573106 | 0.717 | 0.027 | 0.000000e+00 |
| Thbd | 1.9078395 | 0.882 | 0.118 | 0.000000e+00 |
| Cdh5 | 1.8601120 | 0.966 | 0.029 | 0.000000e+00 |
| Adgrf5 | 1.8066178 | 0.940 | 0.050 | 0.000000e+00 |
| Id1 | 1.7850078 | 0.812 | 0.168 | 4.762163e-249 |
| Kdr | 1.7653330 | 0.860 | 0.018 | 0.000000e+00 |

## F

#### Sympathetic neuron Markers

| Gene | Log2Fold change | Percentage 1 | Percentage 2 | Adjusted P Value |
|---|---|---|---|---|
| Snhg11 | 3.2061663 | 0.990 | 0.046 | 0.000000e+00 |
| Insrr | 1.7951871 | 0.997 | 0.062 | 0.000000e+00 |
| Smpd3 | 0.8982419 | 0.895 | 0.085 | 0.000000e+00 |
| Cacna1b | 1.1997503 | 0.978 | 0.075 | 0.000000e+00 |
| Spock3 | 0.7479784 | 0.762 | 0.078 | 8.747048e-264 |
| Gpr22 | 0.5700290 | 0.790 | 0.054 | 0.000000e+00 |
| Tmem59l | 0.7677566 | 0.787 | 0.072 | 0.000000e+00 |
| March11 | 0.6499392 | 0.857 | 0.064 | 0.000000e+00 |
| Sgsm1 | 0.9615743 | 0.956 | 0.057 | 0.000000e+00 |
| Atp2b2 | 0.8224142 | 0.908 | 0.061 | 0.000000e+00 |
| Ptchd1 | 0.8112243 | 0.838 | 0.063 | 0.000000e+00 |
| Dmkn | 0.6906363 | 0.698 | 0.054 | 4.118108e-285 |
| Eml5 | 1.0585886 | 0.911 | 0.036 | 0.000000e+00 |
| Gria2 | 1.2867993 | 0.962 | 0.054 | 0.000000e+00 |
| Arfgef3 | 0.6022328 | 0.883 | 0.036 | 0.000000e+00 |
| Slc27a6 | 0.4985297 | 0.581 | 0.034 | 2.999406e-273 |
| Slc7a14 | 0.6440851 | 0.825 | 0.064 | 0.000000e+00 |
| B3galt1 | 0.6604370 | 0.854 | 0.050 | 0.000000e+00 |
| Plppr5 | 0.5191926 | 0.794 | 0.037 | 0.000000e+00 |
| Shisal1 | 0.4718486 | 0.816 | 0.044 | 0.000000e+00 |

S6

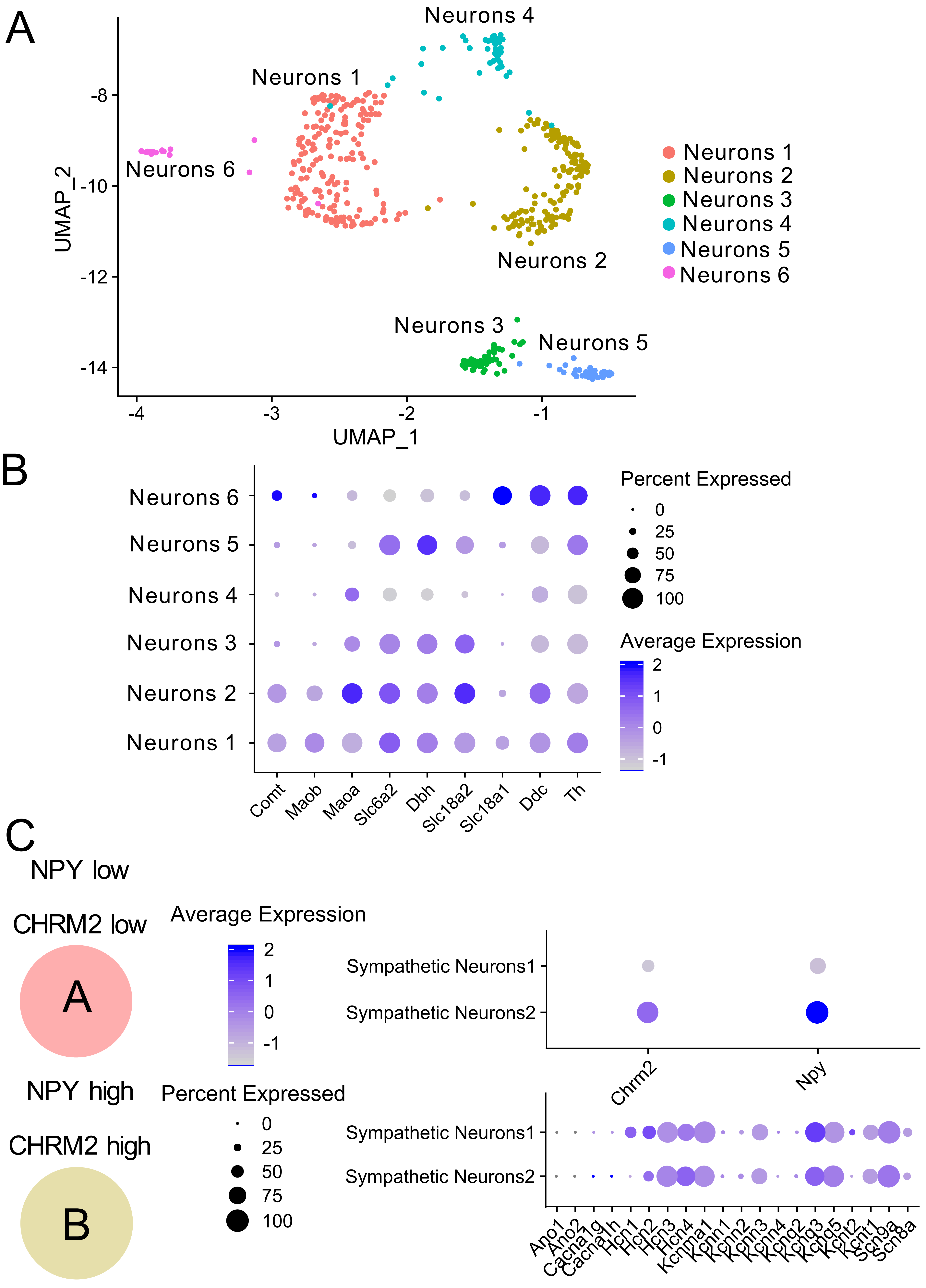
A
Neurons 4
Neurons 1
Neurons 6
Neurons 2
Neurons 3
Neurons 5
-8
-10
-12
-14
UMAP_2
-4
-3
-2
-1
UMAP_1
Neurons 1
Neurons 2
Neurons 3
Neurons 4
Neurons 5
Neurons 6
B
Neurons 6
Neurons 5
Neurons 4
Neurons 3
Neurons 2
Neurons 1
Comt
Maob
Maoa
Slc6a2
Dbh
Slc18a2
Slc18a1
Ddc
Th
Percent Expressed
0
25
50
75
100
Average Expression
2
1
0
-1
C
NPY low
CHRM2 low
A
NPY high
CHRM2 high
B
Average Expression
2
1
0
-1
Percent Expressed
0
25
50
75
100
Sympathetic Neurons1
Sympathetic Neurons2
Chrm2
Npy
Sympathetic Neurons1
Sympathetic Neurons2
Ano1
Ano2
Cacna1g
Cacna1h
Hcn1
Hcn2
Hcn3
Hcn4
Kcnma1
Kcnn1
Kcnn2
Kcnn3
Kcnn4
Kcnq2
Kcnq3
Kcnq5
Kcnt2
Kcnt1
Scn9a
Scn8a

# S7

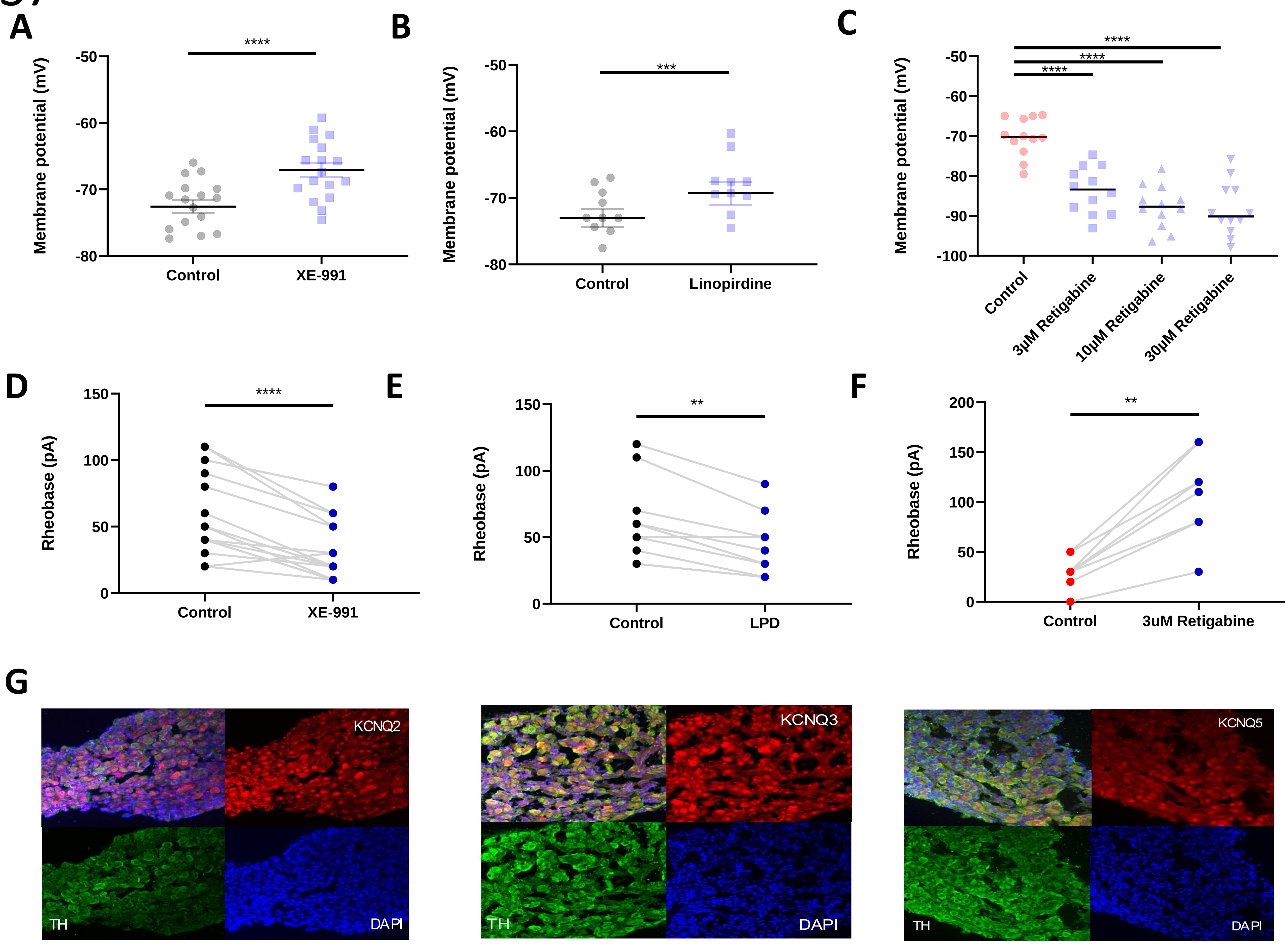
A
B
C
D
E
F
G
Membrane potential (mV)
-50
-60
-70
-80
-90
-100
****
***
Control
XE-991
Linopirdine
3μM Retigabine
10μM Retigabine
30μM Retigabine
Rheobase (pA)
0
50
100
150
200
**
LPD
3uM Retigabine
KCNQ2
KCNQ3
KCNQ5
TH
DAPI

# S8

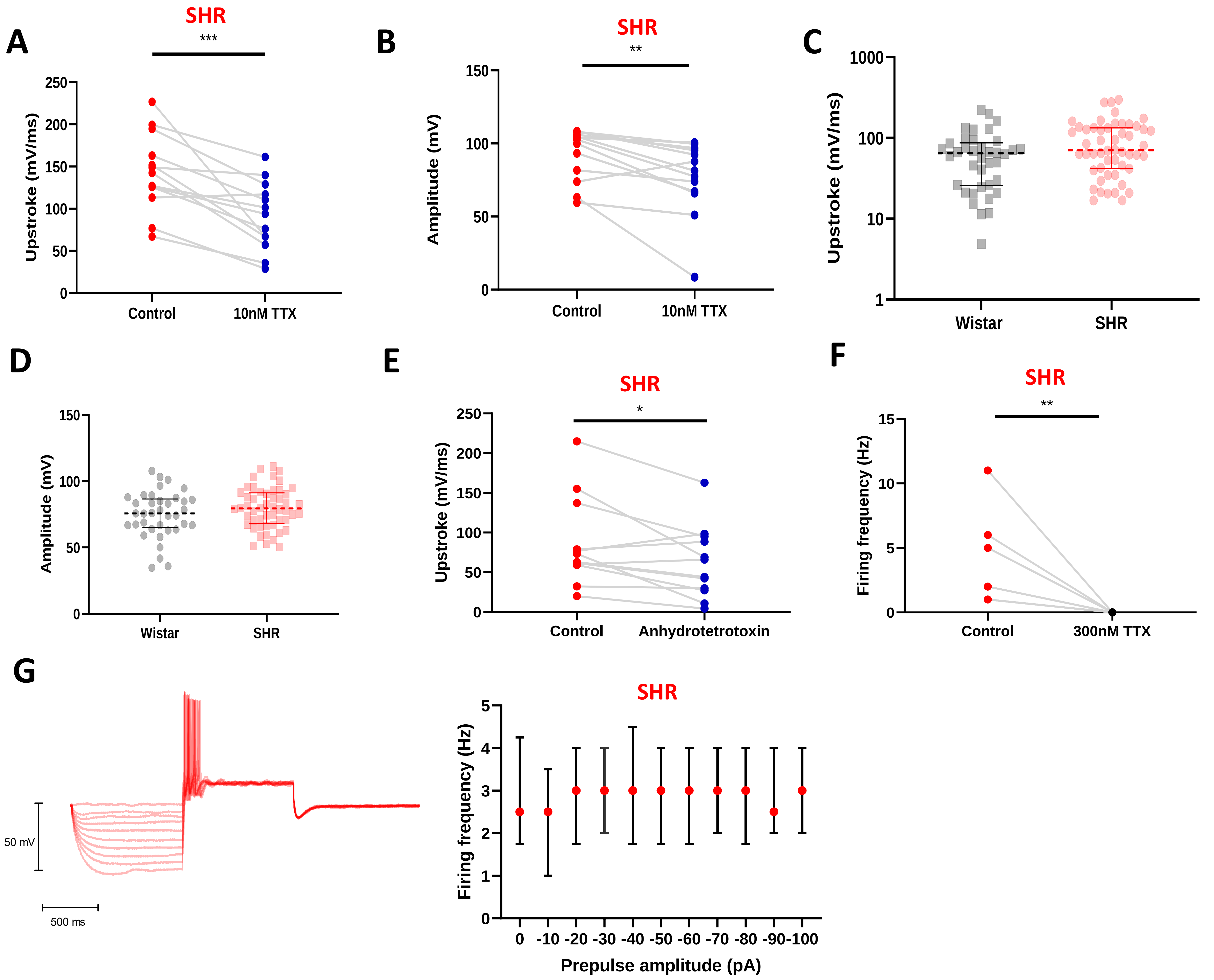

A
SHR
***
Upstroke (mV/ms)
250
200
150
100
50
0
Control
10nM TTX
B
SHR
**
Amplitude (mV)
150
100
50
0
Control
10nM TTX
C
Upstroke (mV/ms)
1000
100
10
1
Wistar
SHR
D
Amplitude (mV)
150
100
50
0
Wistar
SHR
E
SHR
*
Upstroke (mV/ms)
250
200
150
100
50
0
Control
Anhydrotetrotoxin
F
SHR
**
Firing frequency (Hz)
15
10
5
0
Control
300nM TTX
G
50 mV
500 ms
SHR
Firing frequency (Hz)
5
4
3
2
1
0
0 -10 -20 -30 -40 -50 -60 -70 -80 -90 -100
Prepulse amplitude (pA)

# S9

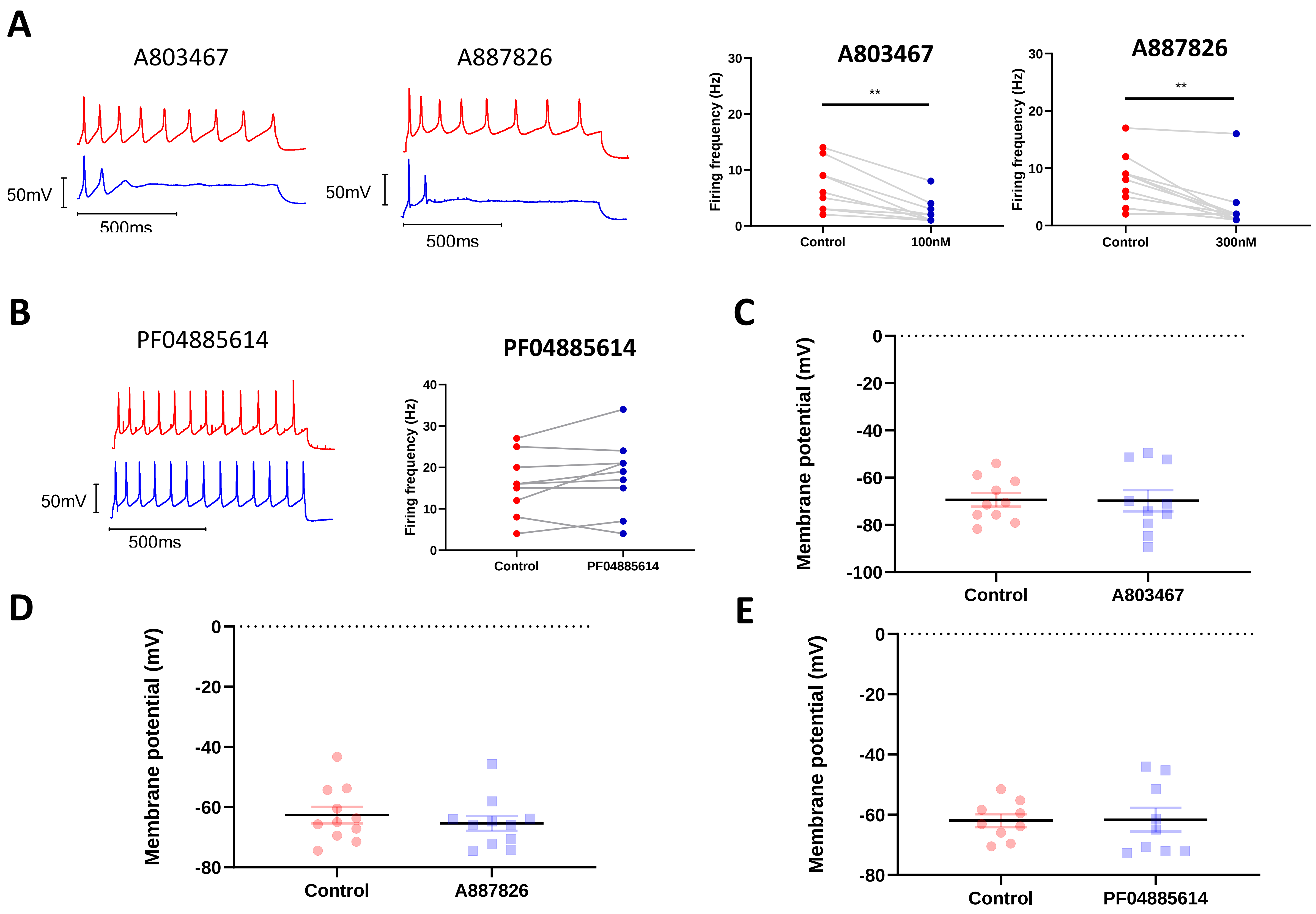
A
A803467
A887826
50mV
500ms
50mV
500ms
A803467
Firing frequency (Hz)
**
Control
100nM
A887826
Firing frequency (Hz)
**
Control
300nM
B
PF04885614
50mV
500ms
PF04885614
Firing frequency (Hz)
Control
PF04885614
C
Membrane potential (mV)
Control
A803467
D
Membrane potential (mV)
Control
A887826
E
Membrane potential (mV)
Control
PF04885614

# S10

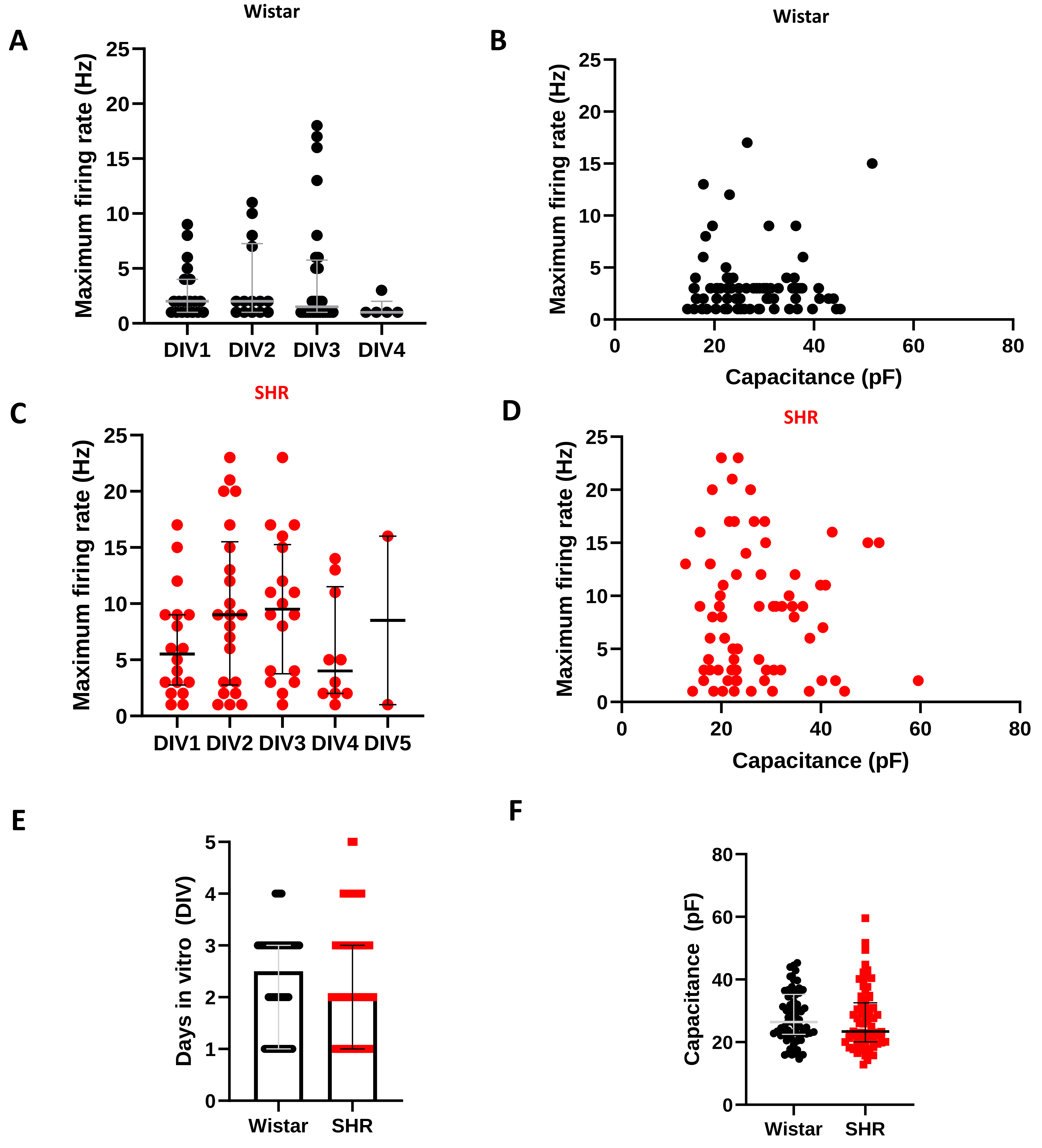

A
Wistar
Maximum firing rate (Hz)
0
5
10
15
20
25
DIV1
DIV2
DIV3
DIV4
B
Wistar
Maximum firing rate (Hz)
Capacitance (pF)
0
20
40
60
80
C
SHR
Maximum firing rate (Hz)
DIV1
DIV2
DIV3
DIV4
DIV5
D
SHR
Maximum firing rate (Hz)
Capacitance (pF)
E
Days in vitro (DIV)
0
1
2
3
4
5
Wistar
SHR
F
Capacitance (pF)
0
20
40
60
80
Wistar
SHR
